# Heliorhodopsin evolution is driven by photosensory promiscuity in monoderms

**DOI:** 10.1101/2021.02.01.429150

**Authors:** Paul-Adrian Bulzu, Vinicius Silva Kavagutti, Maria-Cecilia Chiriac, Charlotte D. Vavourakis, Keiichi Inoue, Hideki Kandori, Adrian-Stefan Andrei, Rohit Ghai

## Abstract

The ability to harness Sun’s electromagnetic radiation by channeling it into high-energy phosphate bonds empowered microorganisms to tap into a cheap and inexhaustible source of energy. Life’s billion-years history of metabolic innovations led to the emergence of only two biological complexes capable of harvesting light: one based on rhodopsins and the other on (bacterio)chlorophyll. Rhodopsins encompass the most diverse and abundant photoactive proteins on Earth and were until recently canonically split between type-1 (microbial rhodopsins) and type-2 (animal rhodopsins) families. Unexpectedly, the long-lived type-1/type-2 dichotomy was recently amended through the discovery of heliorhodopsins (HeRs) (Pushkarev et al. 2018), a novel and exotic family of rhodopsins (i.e. type-3) that evaded recognition in our current homology-driven scrutiny of life’s genomic milieu. Here, we bring to resolution the debated monoderm/diderm occurrence patterns by conclusively showing that HeR distribution is restricted to monoderms. Furthermore, through investigating protein domain fusions, contextual genomic information, and gene co-expression data we show that HeRs likely function as generalised light-dependent switches involved in the mitigation of light-induced oxidative stress and metabolic circuitry regulation. We reason that HeR’s ability to function as sensory rhodopsins is corroborated by their photocycle dynamics (Pushkarev et al. 2018) and that their presence and function in monoderms is likely connected to the increased sensitivity to light-induced damage of these organisms (Maclean et al. 2009).

---

Type-1 and −2 rhodopsins families share a similar topological conformation and little or no sequence similarity among each other. Despite dissimilarities in function, structure and phylogeny, type-1 and −2 rhodopsins have a similar membrane orientation with their N-terminus being situated in the extracellular space. Identified during a functional metagenomics screen and characterised by low sequence similarity when compared to type-1 rhodopsins, HeRs attracted increasing research interest due to their peculiar membrane orientation (i.e. N-terminus in the cytoplasm and the C-terminus in the extracellular space)(Pushkarev et al. 2018), unusual protein structure (Kovalev et al. 2020) and controversial taxonomic distribution (Flores-Uribe, Hevroni, and Ghai 2019). While electrophysiological (Pushkarev et al. 2018), physicochemical (Tanaka et al. 2020) and structural (Shihoya et al. 2019; Kovalev et al. 2020) studies achieved great progress in elucidating a series of characteristics ranging from photocycle length (indicating no pumping activity) to detailed protein organization, they provide no data regarding the biological function of HeRs. Moreover, polarized opinions regarding the putative ecological role and taxonomic distribution of HeR-encoding organisms (Flores-Uribe, Hevroni, and Ghai 2019; Kovalev et al. 2020) call for the use of novel approaches in establishing HeR functionality. This work draws its essence from the tenet that functionally linked genes within prokaryotes are co-regulated, and thus occur close to each other (Aravind 2000; Huynen et al. 2000). Within this framework, the functions of uncharacterised genes (i.e. HeRs) can be inferred from their genomic surroundings. Here we couple HeR’s distributional patterns with contextual genomic information involving protein domain fusions and operon organization, and gene expression data to shed light on HeRs functionality.

Previous assessments of taxonomic distribution of HeRs reported conflicting data regarding their presence in monoderm (Flores-Uribe, Hevroni, and Ghai 2019) and diderm (Kovalev et al. 2020) prokaryotes. In order to accurately map HeR taxonomic distribution we used the GTDB database (release 89), since it contains a wide-range of high-quality genomes derived from isolated strains and environmental metagenome-assembled genomes, classified within a robust phylogenomic framework (Parks et al. 2020). By scanning 24,706 genomes, we identified 450 *bona fide* HeR sequences (topology: C-terminal inside and N-terminal outside, seven transmembrane helices and a SxxxK motif in helix 7; Supplementary Table S1) spanned across 17 phyla (out of 151; Supplementary Table S2). In order to assign HeR-containing genomes to either monoderm or diderm categories, we employed a set of 27 manually curated protein domain markers that are expected to be restricted to organisms possessing double-membrane cellular envelopes (i.e. diderms) (Taib et al. 2020). While most analyses were expected to be influenced by varying levels of genome completeness, we found that a conservative criterion of presence of at least ten marker domains singled out all diderm lineages (i.e. Negativicutes, Halanaerobiales and Limnochondria) (Taib et al. 2020; Megrian et al. 2020) within the larger monoderm phylum Firmicutes, apart from correctly identifying other well-known diderms. Except for three genomes (one each belonging to Myxococcota, Spirochaetota and Dictyoglomota phyla), all other HeR occurrences were restricted to monoderms (Supplementary Table S2). Examination of the HeR-encoding Myxoccoccota contig by querying its predicted proteins against the RefSeq and GTDB databases revealed it to be an actinobacterial contaminant. The *Spirochaeta* genome was incomplete (60% estimated completeness) and only encoded for two outer membrane marker genes, making any inferences regarding its affiliation to monoderm or diderm bacteria impossible. However, we could not rule out that this genome could belong to a member lacking lipopolysaccharides (LPS) (Taib et al. 2020). The Dictyoglomota genome belongs to an isolate, and despite its high completeness, it encodes only five markers. Combined with the notion that Dictyoglomota are known to have atypical membrane architectures (Saiki et al. 1985), the presence of only five markers points towards the absence of a classical diderm cell envelope. Apart from these exceptions, all other HeR-encoding genomes are monoderm and, at least within this collection, we find no strong evidence of HeRs being present in any organism that is conclusively diderm. We also identified HeRs in several assembled metagenomes and metatranscriptomes (see Methods). For improved resolution of taxonomic origin, we considered only contigs of at least 5 Kb in length (n = 1,340 from metagenomes and n = 4 from metatranscriptomes). Following a strict approach to taxonomy assignment (i.e. at least 60 % genes giving best-hits to the same phylum and not just majority-rule), we could designate a phylum for most HeRs. Without any exception, we found that all the contigs that received robust taxonomic classification (n = 1,319) belonged to known monoderm phyla (Supplementary Table S3).

Domain fusions with rhodopsins are recently providing novel insights into the diverse functional couplings that enhance the utility of a light sensor, e.g. the case of a phosphodiesterase domain fused with a type-1 rhodopsin (Ikuta et al. 2020). As far as we are aware, no domain fusions have been described for HeRs yet. In our search for such domain fusions that may shed light on HeR functionality, the MORN repeat (Membrane Occupation and Recognition Nexus, PF02493) was found in multiple copies (typically 3) at the cytoplasmic N-terminus of some HeRs (n = 36). A tentative 3D model for a representative MORN-HeR could be generated and is shown in Figure 1A. These MORN-HeR sequences were phylogenetically restricted to two environmental branches of MAGs recovered from haloalkaline sediments that affiliate to the family *Syntrophomonadaceae* (phylum Firmicutes) (Timmers et al. 2018; Vavourakis et al. 2018, 2019) (Supplementary Figure 1). The prototypic MORN repeat, consisting of 14 amino acids with the consensus sequence YEGEWxNGKxHGYG, was first described in 2000 (Takeshima et al. 2000) from junctophilins present in skeletal muscle and later recognized to be ubiquitous in both eukaryotes and prokaryotes (El-Gebali et al. 2019). This conserved signature can be seen in the alignment of MORN-repeats fused to HeRs (Supplementary Figure 2). MORN-repeats have been shown to bind to phospholipids (Im et al. 2007; Ma et al. 2006), promoting stable interactions with plasma membranes (Takeshima et al. 2000) and also function as protein-protein interaction modules involved in di- and oligomerization (Sajko et al. 2020). They are expected to be intracellular and provide a large putative interaction surface (either with other MORN-HeRs or other proteins). A widespread adaptation of bacteria to alkaline environmental conditions is the increased fluidity of their plasma membranes achieved by the incorporation of branched-chain and unsaturated fatty acids which ultimately influences the configuration and activity of membrane integral proteins such as ATP synthases and various transporters (Kanno et al. 2015). Microbial rhodopsins typically associate as oligomers in vivo, which is also the case with heliorhodopsins that are known to form dimers (Shibata et al. 2018; Shihoya et al. 2019). The presence of MORN-repeats in HeRs exclusively within extreme haloalkaliphilic bacteria (class Dethiobacteria) may be accounted for via their potential role in stabilizing HeR dimers in conditions of increased membrane fluidity (Supplementary Figure 4). Another possibility would be the interaction of MORN-repeats with other MORN-repeat containing proteins encoded in these MAGs. We could indeed identify multiple MORN-protein domain fusions co-occurring in genomes of analysed Dethiobacteria (Supplementary Figures 1 and 3; Supplementary Table S15). Even though the nature of interactions amongst these proteins with intracellular MORN-repeats is unclear, they raise the possibility that MORN-repeats act as downstream transducers of conformational changes occuring in HeRs. Such tandem repeat structures may function as versatile target recognition sites capable of binding not only small molecules like nucleotides but also peptides and larger proteins (Kajava 2012). If true, this would render HeRs as sensory rhodopsins. In support of this, we found several genes in close proximity to MORN-HeRs encoding signature protein domains (e.g. PAS, HisKA, HATPase_c) that are known to be involved in histidine kinase signalling (Aravind, Iyer, and Anantharaman 2010) (Figure 2A).

**Figure 1.**
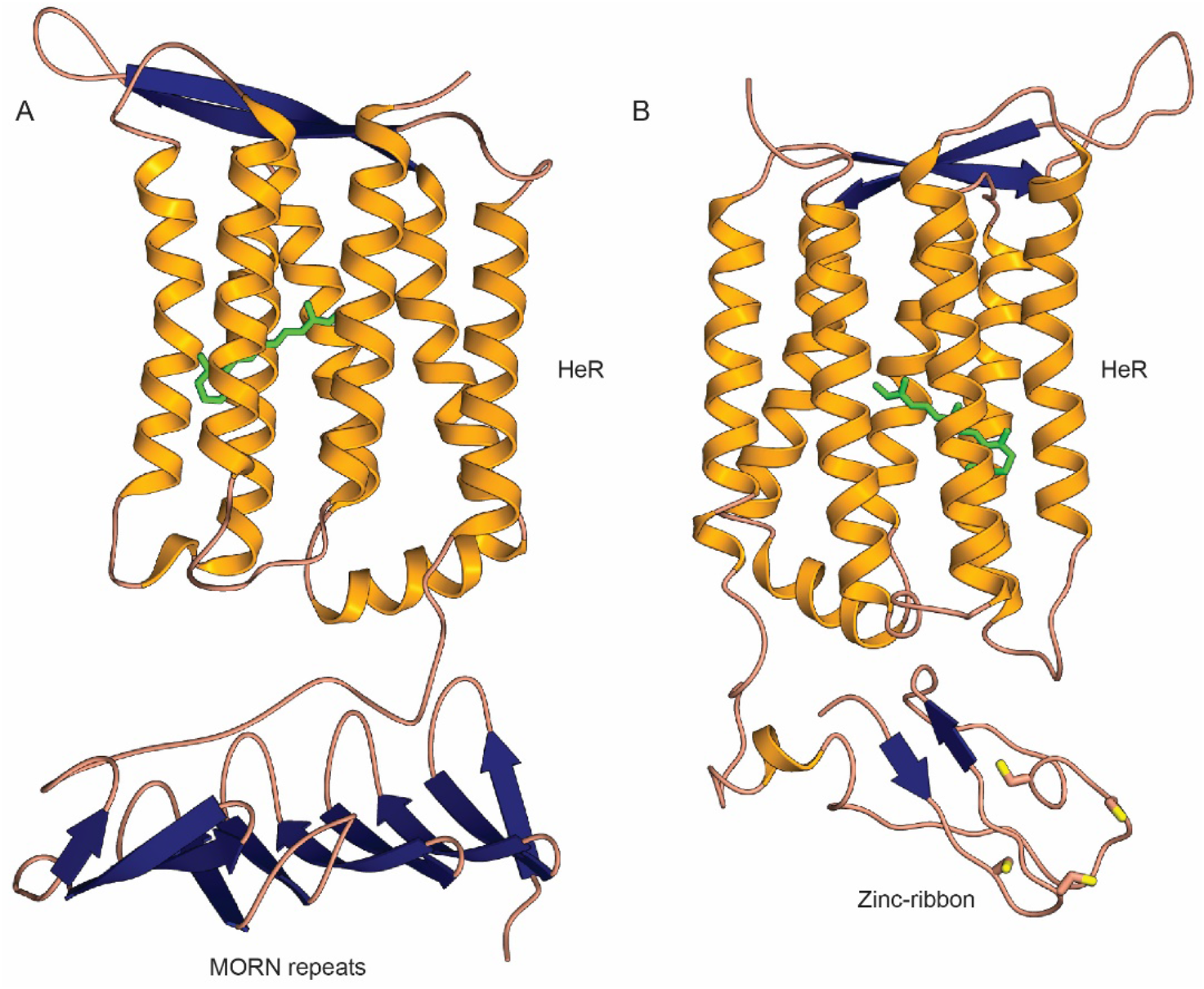
Modelled three-dimensional (3D) structures of MORN-HeR and Znf-HeR protein domain fusions. (A) 3D model of a heliorhdodopsin (HeR) containing three N-terminal MORN domain repeats. (B) 3D model of a HeR containing an N-terminal Zn ribbon motif. Both models are oriented with the extracellular side up and intracellular side down. Retinal is coloured green and cysteine residues are depicted with yellow-topped orange sticks.

**Figure 2.**
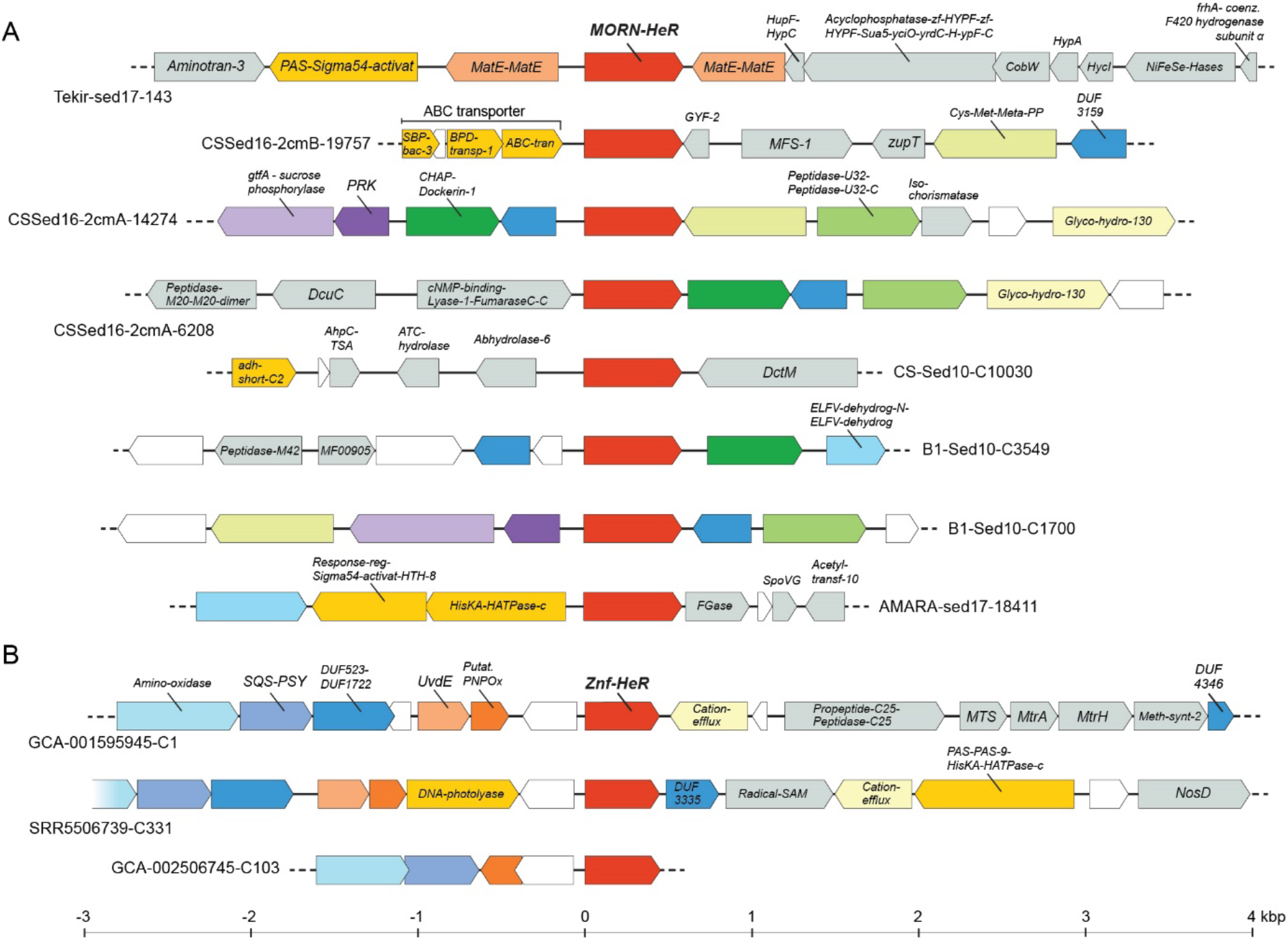
Genomic context of HeR-protein domain fusion genes. A) Representative MORN-HeR encoding contigs identified in strictly anaerobic Firmicutes. B) Contigs encoding Znf-HeR domain fusions. Neighbouring genes were depicted within an interval spanning ~ 7 kb, centered on HeR. Genes occurring only once within the considered intervals are coloured grey; genes encoding HisKA, PAS, regulatory domains, as well as other discussed HeR neighbours are depicted bright yellow. Homologous genes occurring multiple times found within each category of HeR-protein fusion contigs are depicted using matching colours. Hypothetical genes are white.

As no other obvious domains were found to be fused with HeRs using standard profile searches, we examined all N and C-terminal extensions as well as loops longer than 50 aa by performing more sensitive profile-profile searches using HHPred (Zimmermann et al. 2018). We found at least ten N-terminal extensions of HeRs (ntv1-ntv10), 22 variants of ECL1 (extracellular loop 1), a single type of loop extension for ICL2 (intracellular loop 2) and three variants of ICL3 (intracellular loop 3). A complete listing of all alignments and summary results of HHpred can be found in Supplementary Table S8. Remarkably, we found significant matches in a set of six sequences (all originating from Thermoplasmatales archaea) to zinc ribbon proteins (Pfam domain zinc_ribbon_4) at the N-terminus of some heliorhodopsins (these extensions are termed N-terminal variant 1 or ntv1, Supplementary Table S8). Zinc ribbons belong to the larger family of Zinc-finger domains (Krishna et al., 2003). A CxxC-17x-CxxC was found in this region that likely coordinates a metal (e.g., zinc or iron). These CxxC_CxC type motifs are common to a wider family of Zinc finger-like proteins that were initially found to bind to DNA and later shown to be capable of binding to RNAs, proteins and small molecules (Krishna, Majumdar, and Grishin 2003). Similar motifs are also seen in Rubredoxins and Cys_rich_KTR domains. We term these fused ntv1 protein variants as Znf-HeRs (Zinc finger Heliorhodopins). A modelled structure for a representative Znf-HeR is shown in Figure 1. In one contig encoding a Znf-HeR we identified a histidine kinase that could be functionally linked (Figure 2B). Notably, most identified Znf-HeRs are flanked by genes known to be triggered by light exposure and play key roles in photoprotection (i.e. carotenoid biosynthesis genes Lycopene cyclase, phytoene desaturase – Amino-oxidase, squalene/phytoene synthase – SQS-PSY) and UV-induced DNA damage repair (DNA photolyases, UV-DNA damage endonucleases – UvdE) (Rastogi et al. 2010; Yatsunami et al. 2014). Recent research showed that HeRs from Thermoplasmatales archaea (*7a*HeR) and uncultured freshwater Actinobacteria (48C12) (for which the structure is resolved and lacks the ntv1 extension) might bind zinc (Hashimoto et al. 2020). As the zinc binding site could not be precisely identified it was suggested that it could be located in the cytoplasmic part, and responsible for modifying the function of HeR. Our discovery of Znf-HeRs offers additional, more direct indications of the role of zinc in the possible downstream signalling by HeRs.

Given the large number of long contigs encoding HeRs (from genomes and metagenomes), we sought to identify candidate genes that could be transcribed together with HeRs (in the same operon). We used the following strict criteria for obtaining such genes 1) the intergenic distance between such a gene and the HeR must be less than 10 bp, and 2) the gene must be located on the same strand. A number of interesting candidates emerged in this analysis with the most frequent ones being summarized in Figure 3 (a complete table can be found in Supplementary Table S9). We identified multiple instances in which genes with Glutaredoxin and GSHPx PFAM domains were found adjacent to HeRs (n = 31). Glutaredoxins are small redox proteins with active disulphide bonds that utilize reduced glutathione as an electron donor to catalyze thiol-disulphide exchange reactions. They are involved in a wide variety of critical cellular processes like the maintenance of cellular redox state, iron and redox-sensing, and biosynthesis of iron-sulphur clusters (Lillig, Berndt, and Holmgren 2008; Rouhier et al. 2010). Glutathione is also used by glutathione peroxidase (GSHPx) to reduce hydrogen peroxide and peroxide radicals i.e. as an anti-oxidative stress protection system (Bhabak and Mugesh 2010). Additionally, there are also instances where Glutaredoxin and genes containing Glyoxalase_2 domains may be co-transcribed with HeRs. Glyoxalases, in concert with glutaredoxins, are critical for detoxification of methylglyoxal, a toxic byproduct of glycolysis (Ferguson et al. 1998). Moreover, adjacent to HeRs we find at least three instances where a catalase gene is also present (in Actinobacteria; see Supplementary Figures S10–S11). Collectively, these observations suggest a role for HeRs in oxidative stress mitigation. In one case, we found a gene encoding the DICT domain (Figure 3) which is frequently associated to GGDEF, EAL, HD-GYP, STAS, and two-component system histidine kinases. Notably, it has been predicted to have a role in light response (Aravind, Iyer, and Anantharaman 2010).

**Figure 3.**
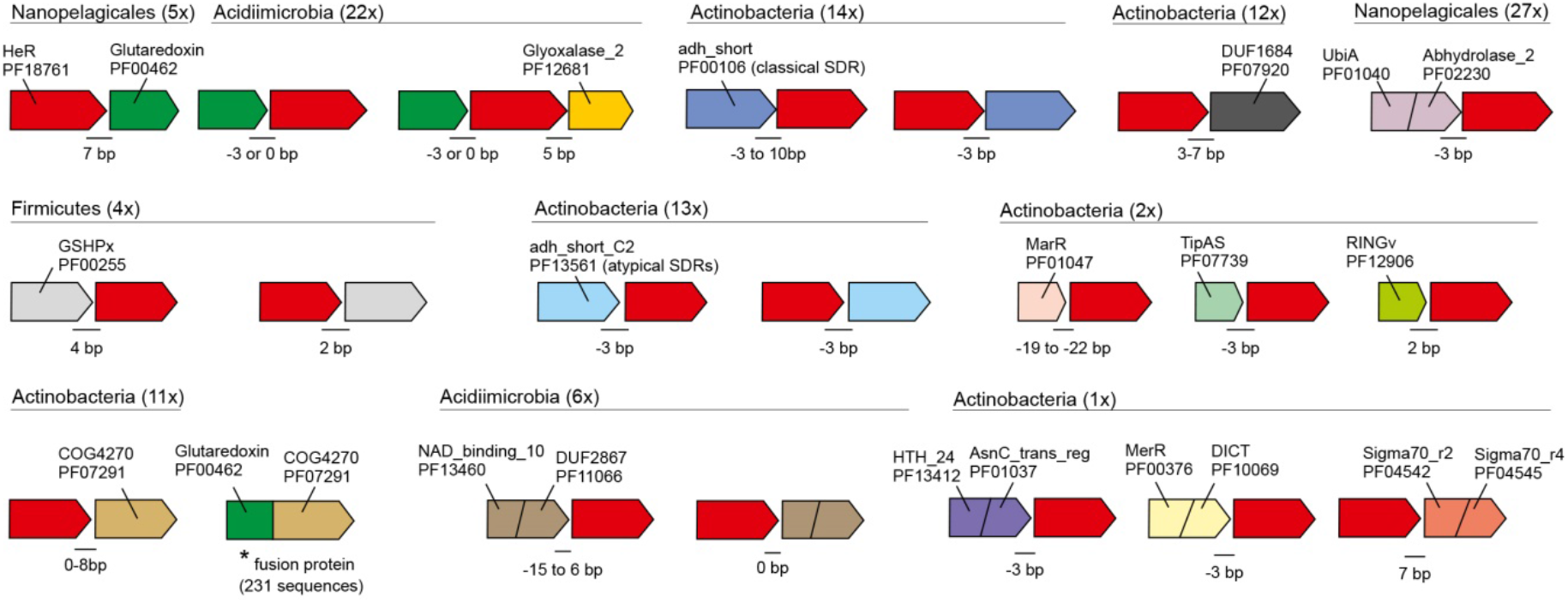
Schematic representation of genes that may be transcriptionally linked to HeRs. Taxonomic categories and number of occurrences are shown at the top of each putative operon. Intergenic distances (in bp) are indicated at gene junctions. Negative distance values indicate overlapping genes. Pfam or COG identifiers are used to represent domain architectures. A star (*) indicates a fused gene (two domains: Glutaredoxin and COG4270) found in at least 473 genomes from GTDB and 231 unique sequences in UniProt suggesting a functional linkage of COG4270 with Glutaredoxin.

Although we assembled contigs encoding HeRs from previously published metatranscriptomes, the lack of strand-specific transcriptomes hampered any clear conclusions on whether or not genes adjacent to HeRs are indeed co-transcribed, leaving open the possibility that they might simply be artefacts of assembly (Zhao et al. 2015). In order to gather more definitive evidence for co-transcription we performed strand-specific metatranscriptome sequencing for a freshwater sample (see Methods). We recovered six HeR-encoding transcripts that were > 1 kb in length. All these transcripts are predicted to originate from highly abundant freshwater Actinobacteria with streamlined genomes (four transcripts from *“Ca. Planktophila”* and two from *“Ca. Nanopelagicus”*) (Supplementary Table S12) (Neuenschwander et al. 2018). Overall, there are three types of transcripts based upon gene content: class1 – encoding Glutamine synthetase catalytic subunit and NAD+ synthetase; class2 – encoding a hydrolase, a peptidase and a DUF393 domain containing protein, and class3 – encoding glucose/sorbosone dehydrogenase (GSDH) (Figure 4B and Supplementary Table S12). A common theme for glutamine synthetase and NAD+ synthetase is that both utilize ammonia and ATP to produce glutamine and NAD+ respectively. Moreover, some NAD+ synthetases may be glutamine dependent (Resto, Yaffe, and Gerratana 2009). Glutamine synthetase in particular is a key enzyme for nitrogen metabolism in prokaryotes at large (García-Domínguez, Reyes, and Florencio 1999). For hydrolases and peptidases, the function prediction is somewhat broad. Glucose/sorbosone dehydrogenase catalyses the production of gluconolactone from glucose (Oubrie et al. 1999). Therefore, it appears that all six HeRs are generally co-transcribed with genes involved in nitrogen assimilation and degradation/assimilation of sugars and peptides. This would suggest that these processes are also influenced by light, with such a link between light-dependent increase in sugar uptake and metabolic activity being recently proposed in non-phototrophic Actinobacteria (Maresca et al. 2019). Light also triggers photosynthetic activity, increasing availability of sugars and other nutrients (e.g. glutamine and ammonia) for heterotrophs. In this vein, a link between a light sensing mechanism, e.g. via heliorhodopsins, may lead to elevated metabolic activity.

**Figure 4.**
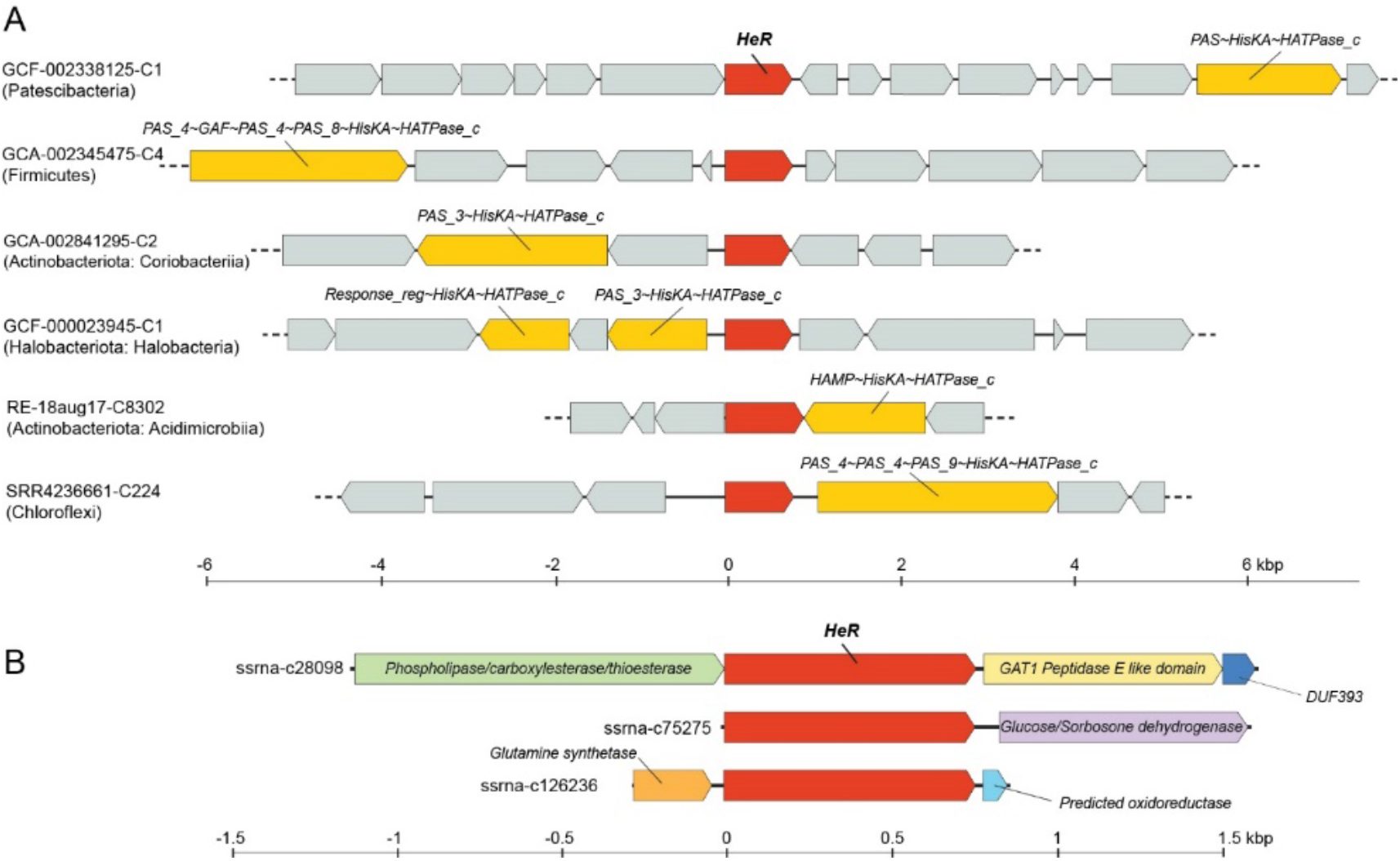
A) Genes encoding HisKA domain signalling proteins identified in the proximity of HeR genes from diverse phyla. All genes containing HisKA domains are coloured bright yellow, HeRs are shown in red, and all other genes in grey. B) Transcripts obtained by strand-specific metatranscriptomics from freshwater encoding genes co-expressed with HeR.

In a previous study, histidine kinases were deemed absent in the vicinity of HeRs (Kovalev et al. 2020). Given that our initial analyses predicted a sensory function, we examined genomic regions spanning 10 kb up- and downstream of HeRs. Already in the case of MORN-HeRs and Znf-HeRs we observed histidine kinase signalling components in close proximity to them (Figure 2). In our search we detected multiple instances of histidine kinases (HisKA) fused with PAS, GAF, MCP_Signal, HAMP or HATPase_c domains in the gene neighbourhoods of HeRs in distinct phyla (e.g. Actinobacteria, Chloroflexi, Patescibacteria, Firmicutes, Dictyoglomota, Thermoplasmatota) (Figure 4B; more details in Supplementary Figures S5–S14). Moreover, in many cases multiple response regulator genes were present in the same regions (Pfam domains Response_reg, Trans_reg_C).

Less frequently, GGDEF and EAL domains, usually associated with bacterial signalling proteins, were also present. Using overrepresentation analysis (Shmakov et al. 2018), we found that the occurrence of two-component system protein domains in the vicinity of HeRs is statistically significant (see Methods and Supplementary Table S11). In addition to these two-component system proteins, the same regions also appear enriched in redox proteins (e.g. thioredoxin, peroxidase, catalase). The close association of two-component systems, genes involved in oxidative stress mitigation and HeRs points towards a functional interaction.

## Conclusions

In conclusion, contextual genomic information shows that monoderm prokaryotes use HeRs in multiple mechanisms for the activation of downstream metabolic pathways post light sensing. Furthermore, we offer tantalizing clues regarding the involvement of HeRs in multiple cellular processes and add new lines of inquiry for the primary role of HeRs in light signal transduction. Additional support for the role of HeRs in light sensing is inferred from the frequent association of HeRs with classical histidine kinases and associated protein domains in multiple phyla. Furthermore, multiple types of N-terminal domain fusions found in specific subfamilies of HeRs (i.e. MORN domains in haloalkaliphilic Firmicutes and Zinc ribbon type domains in Thermoplasmatales archaea) point to possible downstream signalling which may be effected by recruitment of additional, as yet unknown, partner proteins.

We further propose a critical role for HeRs in protecting monoderm cells from light-induced oxidative damage. In this sense, we observed a close association and probable transcriptional linkage of HeRs to glyoxylases and glutaredoxins (sometimes seen as overlapping genes). Given that light can induce the uptake and metabolism of sugars, as previously discussed for certain Actinobacteria (Maresca et al. 2019), it is expected that increased sugar availability resulting from photosynthesis leads to increased glycolytic activity in heterotrophic bacteria. Glycolysis also produces small amounts of toxic methylglyoxal that can be neutralized by the combined action of glyoxylases and glutaredoxins. In this sense, it appears that at least in some Actinobacteria, glyoxylases and glutaredoxins may be transcribed together with HeRs, but how the transcription is controlled remains unclear. Additional evidence of transcriptional linkages of HeRs to proteins like peroxiredoxin and catalase also imply a light-dependent activation, boosting the cellular response to light induced oxidative damage which may be critical for both aerobes and anaerobes. Evidence from strand-specific HeR transcripts originating from freshwater Actinobacteria suggests the further involvement of HeRs in nitrogen and sugar metabolism via glutamate synthase, NAD+ synthases and glucose/sorbosone dehydrogenases in these organisms.

Overall, the picture that emerges (at least for some organisms) is one of HeR’s role in responding to light and transmitting the signal via histidine kinases. Downstream processes that are ultimately regulated are diverse, including possible roles for HeRs in the mitigation of light-induced oxidative damage and in the regulation of nitrogen assimilation and carbohydrate metabolism, processes that may benefit from a light-dependent activation through more efficient utilization of available resources.

Recent work has shown more support for the diderm-first ancestor (Coleman et al. 2020) and given the far broader distribution of type-1 rhodopsins in both mono- and di-derm organisms it appears likely that type-1 rhodopsins emerged prior to HeRs. The very restricted distribution of HeRs to monoderms would support this view as well. Even so, HeRs are not universally present in monoderms and when present, appear to be associated with diverse genes involved in signal transduction, oxidative stress mitigation, nitrogen and glucose metabolism. This would suggest they have been exapted as generalized sensory switches that may allow light-dependent control of metabolic activity in multiple lineages, somewhat similar to type-1 rhodopsins where minor modifications have led to emergence of a wide variety of ion-pumps (Kandori 2020). The frequent distribution of HeRs in aquatic environments (habitats characterised by increased light penetration), where they commonly occur within phylum Actinobacteriota, helps us to explain their monoderm-restricted presence. Abundant freshwater actinobacterial lineages are generally typified by lower GC content (Ghai, McMahon, and Rodriguez-Valera 2012) and increased vulnerability to oxidative stress damage (Kim et al. 2019). This susceptibility is also illustrated by actinobacterial phages that exhibit positive selection towards reactive oxygen species defense mechanisms (Kavagutti et al. 2019). Given the fact that monoderms are generally more sensitive to light-induced damage and corroborated with up-mentioned metabolic implications, we consider that HeRs evolved as sensory switches capable of triggering a fast response against photo-oxidative stress in prokaryotic lineages more sensitive to light.

## Methods

### Metagenomes and metatranscriptomes

We used previously published metagenomics and metatranscriptomics data from freshwaters (Andrei et al. 2019; Kavagutti et al. 2019; Mehrshad et al. 2018), haloalkaline brine and sediment (Vavourakis et al. 2018, 2019), brackish sediments (Bulzu et al. 2019), GEOTRACES cruise (Biller et al. 2018) and TARA expeditions (Salazar et al. 2019). In addition, we downloaded multiple environmental metagenomes (sludge, marine, pond, estuary, etc.) from EBI MGnify (https://www.ebi.ac.uk/metagenomics/) (Mitchell et al. 2020) and assembled them using Megahit v1.2.9 (D. Li et al. 2016). All contigs in this work are named or retain existing names that allow tracing them to their original datasets.

### Sequence search for *bona fide* rhodopsins

Genes were predicted in metagenomics contigs using Prodigal (Hyatt et al. 2010). Candidate rhodopsin sequences were scanned with hmmsearch (Eddy 2011) using PFAM models (PF18761: heliorhodopsin, PF01036: bac_rhodopsin) and only hits with significant e-values (< 1e-3) were retained. Homologs for these sequences were identified by comparison to a known set of rhodopsin sequences (Bulzu et al. 2019) using MMSeqs2 (Hauser, Steinegger, and Söding 2016) and alignments were made using MAFFT-linsi(Katoh and Standley 2013). These alignments were used as input to Polyphobius (Käll, Krogh, and Sonnhammer 2005) for transmembrane helix prediction. Only those sequences that had seven transmembrane helices and either a SxxxK motif (for heliorhodopsins) or DxxxK motif (for proteorhodopsins) in TM7 were retained. In addition, we also screened the entire UniProtKB for HeRs. In total, we accumulated at least 4,1 08 (3,606+502) *bona fide* HeR sequences.

### Taxonomic classification of assembled contigs

Contigs were dereplicated using cd-hit (W. Li and Godzik 2006) (95% sequence identity and 95% coverage). Only contigs ≥ 10 kb were retained for this analysis. A custom protein database was created by predicting and translating genes in all GTDB genomes (release 89) (Parks et al. 2020) using Prodigal (Hyatt et al. 2010). These sequences were supplemented with viral and eukaryotic proteins from UniProtKB (UniProt Consortium 2019). Best-hits against predicted proteins in contigs were obtained using MMSeqs2 (Hauser, Steinegger, and Söding 2016). Taxonomy was assigned to a contig (minimum length 5 kb) only if ≥ 60% of genes in the contig gave best-hits to the same phylum. All contigs that appeared to originate from diderms were cross-checked against NCBI RefSeq (accessed online on 15^th^ December 2020).

### Outer-envelope detection

A set of protein domains found in genes encoding for the outer-envelope (Taib et al. 2020) was further reduced to include only those domains that were found mostly in known diderms. These domains were searched against the predicted proteins in all genomes in GTDB using hmmsearch (e-value < 1e-3). The results are shown in Supplementary Table S13.

### Protein function/structure predictions

Predicted proteins were annotated using TIGRFAMs (Haft, Selengut, and White 2003) and COGs (Galperin et al. 2015). Domain predictions were carried out using the pfam_scan.pl script against the PFAM database (release 32) (El-Gebali et al. 2019). Profile-profile searches were carried out online using the HHPred server (Zimmermann et al. 2018). Additional annotations were added using Interproscan (Mitchell et al. 2019). Protein structure predictions were carried out using the Phyre2 server (Kelley et al. 2015) and structures were visualized with CueMol (http://www.cuemol.org/en/).

### Domains overrepresentation near heliorhodopsin

A subset of high-quality MAGs (n = 240) containing HeR-encoding genes flanked both up- and downstream by a minimum of 10 genes were selected from GTDB (release 89) (Parks et al. 2020). For each genome, the probability of finding any particular domain by chance in a random subset of 20 genes was calculated using the hypergeometric distribution (without replacement) in R with the function *phyper (stats* package) (Johnson, Kemp, and Kotz 2005). In order to account for type I errors arising from multiple comparisons, hypergeometric test P-values were adjusted using the Benjamini-Hochberg procedure (Benjamini and Hochberg 1995). Further, we selected domains located in the proximity of HeR in at least 10% of genomes with low probability (FDR corrected P-value < 0.05). This procedure that was initially employed for the whole GTDB genome collection was repeated for individual phyla containing HeR-encoding genes within at least five genomes.

### Strand-specific freshwater transcriptome sequencing and assembly

Sampling was performed on the 16^th^ of August 2020 at 9:00 in Řimov reservoir, Czech Republic, (48°50′54.4″N, 14°29′16.7″E) using a hand-held vertical Friedinger (2 L) sampler. A total of 20 L of water were collected from a depth of 0.5 m and immediately transported to the laboratory. Serial filtration was carried out by passing water sample through a 20 μm pore size pre-filter mesh followed by a 5 μm pore size PES filter (Sterlitech) and a 0.22 μm pore size PES filter (Sterlitech, USA) using a Masterflex peristaltic pump (Cole-Palmer, USA).

Filtration was done at maximum speed for 15 minutes to limit cell lysis and RNA damage as much as possible. A total volume of 3.7 L was filtered during this time. PES filters (5 μm and 0.22 μm pore sizes) were loaded into cryo-vials pre-filled with 500 μl of DNA/RNA Shield (Zymo Research, USA) and stored at −80°C. RNA was extracted from filters using the Direct-zol RNA MicroPrep (Zymo Research, USA) after they had been previously thawed, partitioned, and subjected to mechanical lysis by bead-beating in ZR BashingBead^™^ Lysis tubes (with 0.1 and 0.5 mm spheres). DNase treatment was performed to remove genomic DNA during RNA extraction as an “in-column” step described in the Direct-zol protocol and was repeated after RNA elution, by using the Ambion Turbo DNA-freeTM Kit (Life Technologies, USA). RNA was quantified using a NanoDrop^®^ ND-1000 UV-Vis spectrophotometer (Thermo Fisher Scientific, USA) and integrity was verified by agarose gel (1 %) electrophoresis. A total of 4.6 μg of RNA from the 0.22 μm pore size filter and 2.6 μg from the 5 μm pore size filter were sent for dUTP-marking based strand-specific metatranscriptomic sequencing at Novogene (www.novogene.com). Following quality control at Novogene, samples were mixed into one single reaction, then subjected to rRNA depletion and used for stranded library preparation. Strand-specificity was achieved by incorporation of dUTPs instead of dTTPs in the second-strand cDNA followed by digestion of dUTPs by uracil-DNA glycosylase to prevent PCR amplification of this strand. Paired-end (PE 150 bp) sequencing was carried out using a Novaseq 6000 platform. A total of 166,213,184 raw sequencing reads, amounting to 24.9 Gb, were produced. *De novo* assembly of metatranscriptomic data was performed using rnaSPAdes v.3.14.1 (Bushmanova et al. 2019) in reverse-forward strand-specific mode (--ss rf) with a custom k-mers list 29, 39, 49, 59, 69, 79, 89, 99, 109, 119, 127. A total of 156,235 hard-filtered transcripts of a minimum length of 1 kb were assembled. Protein coding sequences were predicted *de novo* using Prodigal (Hyatt et al. 2010) in metagenomic mode (-p meta). Protein domains were annotated by scanning with InterProScan(Mitchell et al. 2019) while PFAM (Protein Families)(El-Gebali et al. 2019) domains were identified using the publicly available perl script pfam_scan.pl (ftp://ftp.ebi.ac.uk/pub/databases/Pfam/Tools/). Proteins were scanned locally using HMMER3 (Eddy 2011) against the COGs (Clusters of Orthologous Groups) (Galperin et al. 2015) HMM database (e-value < 1e-5) and the TIGRFAMs (TIGR Families) (Haft, Selengut, and White 2003) HMM collection with trusted score cutoffs. BlastKOALA (Kanehisa, Sato, and Morishima 2016) was used to assign KO identifiers (KO numbers). Annotations for representative transcripts encoding HeR are summarised in Supplementary Table S12.

## Data availability

Sequence data generated in this study have been deposited in the European Nucleotide Archive (ENA) at EMBL-EBI under project accession number PRJEB35770 (run ERR5100021). The derived data that support the findings of this paper, including R code used for statistical analyses, are available in FigShare (https://figshare.com/s/7bb42426f2ad5e891fec). All other relevant data supporting the findings of this study are available within the paper and its supplementary information files.

## Acknowledgements

P.-A.B., V.S.K. and R.G. were supported by the research grant 20-12496X (Grant Agency of the Czech Republic). V.S.K. was additionally supported by the research grant 116/2019/P (Grant Agency of the University of South Bohemia in České Budějovice, 2019-2021). A.-Ş.A. was supported by Ambizione grant PZ00P3_193240 (Swiss National Science Foundation). M.-C. C. was supported by the Program for the Support of Perspective Human Resources (PPLZ), Czech Academy of Sciences (Grant No. L200961953). K. I. was supported by Grants-in-Aid from the Japan Society for the Promotion of Science (JSPS) for Scientific Research (KAKENHI grant Nos. 20K21383 and 20H05758). H. K. was supported by a research grant from the Japanese Ministry of Education, Culture, Sports, Science and Technology (18H03986) and a grant from CREST, Japan Science and Technology Agency (JPMJCR1753).

## Author contributions

R.G. and P.-A.B. designed the study. P.-A.B., A.-Ş.A. and R.G. wrote the manuscript. P.-A.B., R.G., V.S.K, M.-C.C., C.D.V and A.-Ş.A. analysed and interpreted the data. K.I. and H.K. performed rhodopsin structural analyses. All authors commented on and approved the manuscript.

## Supplementary Information

We examined the distribution and co-occurrence of HeRs and type-1 rhodopsins (Supplementary Tables S4-S7) in the GTDB database (release 89), as it has been previously suggested that these proteins tend to coexist within the same organisms (Kovalev et al. 2020). From all 24,706 scanned genomes we identified and retrieved 1,455 *bona fide* type-1 rhodopsin-containing genomes from which 69 (4.74 %) proved to also harbour HeRs. Since 15.33 % from all identified HeRs (n = 450) co-occur with type-1 rhodopsins we consider it as being more an exception than a norm and find no data to sustain a physiological dependency between these two rhodopsin families.

### MORN-protein domain fusions

The bacteria encoding MORN-HeRs were previously predicted to be strict anaerobes (Timmers et al. 2018). Although mostly recovered from sediments, these MAGs also encode other proteins directly or indirectly associated with the presence of light (i.e. bacteriophytochrome COG4251, DNA repair photolyase COG1533, Deoxyribodipyrimidine photolyase COG0415). Therefore, the co-occurrence of strict anaerobiosis and light-dependent components indicates the top sediment layer as their likely habitat. The available 3D structure of MORN-repeats shows a single repeat to consist of short beta-pleated regions folded back upon themselves, creating a flat surface area that expands when the repeats are present in multiple tandem copies (Wilson et al., 2002). The presence of periplasmic proteins with MORN-Big_2 or MORN-Big2-PASTA domains may indicate the extracellular MORN repeats as adaptors between typical sensor domains (i.e. PASTA/Big_2) and the transducer protein kinases acting in the cytoplasm (Supplementary Figure 3). However, as the MORN-repeats that are fused to HeRs in these organisms are intracellular, they could only interact with MORN-repeats on the cytoplasmic side. In one such case, intracellular MORN-repeats were fused to ATPase component of an ABC-type efflux pump (likely involved in drug resistance or Cu2+/Na2 + ion efflux). The other candidate found was a PknB protein kinase-FHA-MORN repeat fusion that was predicted to be in an atypical membrane orientation. In such proteins, dimerization domains (e.g. PASTA) are extracellular (Supplementary Figure 3C) while cytoplasmic protein kinase domains function as part of signal transduction pathways in a wide range of gram-positive bacteria (Kang et al. 2005). The dimerization of PknB is essential for autophosphorylation and activation of the kinases through an allosteric mechanism (Lombana et al. 2010). The predicted reverse orientation (if correct) of this protein renders MORN-mediated interactions with HeR unlikely in these organisms. However, the presence of MORN-repeats in PknB type proteins for which dimerization is essential for function (most likely via MORN-repeats) strengthens the possibility that the MORN-repeats aid in MORN-HeR dimerization as well.

### Various Cysteine-rich motif-containing heliorhodopsins

Another extension found in four sequences (N-terminal variant 2 or ntv2 in Supplementary Table S8) with many more cysteines (n = 10) that did not give any significant hits to known proteins was also identified. The presence of multiple cysteine residues and a conserved tryptophan residue is reminiscent of RINGv finger domains that coordinate two metals. However, RING finger domains also have a highly conserved histidine residue that was not detected. Indeed, we also found at least two instances in freshwater Actinobacteria (*Ca. Planktophila*) where a RINGv domain-containing protein is located right upstream of the heliorhodopsin gene and in the same orientation (see Figure 3).

A third variant (ntv3) with seven cysteines was found (in Thermoplasmatales and in Euryarchaeota) but without conserved histidines or tryptophan. However, it shows broad similarity with ntv3 in presence of conserved prolines and arginines before the cysteine motifs. Apart from the n-terminal extensions motifs, at least three sequences of the intracellular loop (ICL3), were also found to be rich in cysteines (n = 8) which also presented a conserved tryptophan similar to ntv2 described above (intracellular loop 3 variant 1, icl3v1).

Thus, at least 17 sequences presented cysteine-rich motifs either at the N-terminus (ntv1, ntv2 and ntv3) or in the intracellular loop (icl3v1) at the cytoplasmic side of heliorhodopsins suggesting the possibility that these might be transducers of the conformational change in heliorhodopsins upon light excitation. The conserved cysteines in these proteins could bind either iron or zinc and are likely redox-active. While we did find many other types of N-terminal extensions, we were unable to find any significant hits to these even by sensitive sequence searches (Supplementary Table S8).

### Sources of retinal for HeR function

We also found several NAD-dependent short-chain dehydrogenases that might encode for retinol dehydrogenases in the vicinity of HeRs (adh_short in Figure 3, Supplementary Table S9). It has been mentioned before that as HeRs are able to efficiently capture retinal from exogenous sources and that HeR-encoding microbes do not have a retinal biosynthesis pathway (Shihoya et al. 2019).

*De novo* retinal biosynthesis requires five genes that if supplied in-trans to a non-retinal producing microbe may result in functional rhodopsins (Sabehi et al. 2005). The final step of the *de novo* pathway uses a beta-carotene monooxygenase that converts beta-carotene to retinal. However, retinal may also be converted from retinol by the action of retinol dehydrogenases. We used the curated GTDB database to further probe the co-occurrence of HeR and type-1 rhodopsins along with genes for retinal biosynthesis. Of the total of 381 genomes that encoded only HeR, we find that only a single genome encoded all genes necessary for the *de novo* production of retinal, but 213 (55%) also encoded the beta-carotene monooxygenase and 241 genomes (63%) encoded at least one retinol dehydrogenase (Supplementary Table S10). Considering genomes that encoded only type-1 rhodopsins (n = 1,386), 596 (43%) encoded the complete pathway for retinal biosynthesis and additionally 995 (71%) also encoded at least one retinol dehydrogenase. It appears that microbes encoding only HeR mostly lack the complete pathway for *de novo* retinal biosynthesis and that apart from exogenous capture of retinal, conversion from beta-carotene (via beta-carotene monooxygenase) or from retinol (via retinol dehydrogenases) may be at work.

### HeR genomic context

We performed gene context analysis of HeRs by combining the maximum-likelihood phylogenetic tree generated for representative HeR sequences (n = 872) with HeR gene neighbourhood information (Supp. Figure 4; iTOL: https://itol.embl.de/tree/14723125092152021608050562). The resulting tree places most HeR sequences (n = 835) within 19 conspicuous phylogenetic clusters which we further denominate as C1-C19 (see Supp. Figure 15). Among them, Actinobacteriota-encoded HeRs are by far the most numerous (n = 533) accounting for eight well-defined clusters (i.e. C1 −7 and C11) and a small sub-cluster within Patescibacteria (CPR)-dominated C17. Regarding Actinobacteriota, C1 and C3 are represented by order Nanopelagicales, C2 includes chiefly members of Microtrichales (class Acidimicrobiia), C4 comprises Actinomycetales HeRs and C5 includes classes Coriobacteriia and Thermoleophilia. Notably, C5 brings together HeRs recovered mainly from lesser studied sediment habitats including nine Chloroflexota (class Dehalococcoidia) sequences. Predominantly marine C6 includes Acidimicrobiia HeRs from order Microtrichales and other poorly classified representatives from within this class while C7 has both marine and freshwater Microtrichales together with Propionibacteriales genus *Nocardioides*.

Cluster C11 stands out in this analysis due to the high level of evolutionary conservation of both HeRs and their neighbouring genes, bringing together exclusively members of marine Actinobacteria from order *Ca*. Actinomarinales (TMED189). Importantly, in C11 we notice that synteny is only conserved among genes sharing the same orientation as HeR while gene “gains” and/or “losses” occur only in the opposite orientation. Despite very high phylogenetic relatedness within C11 and therefore the unsurprisingly similar gene context amongst its members, the differences between (+) and (-) strand feature conservation (relative to HeR orientation) indicate a potentially relevant transcriptional unit comprised of genes: afuA – iron(III) transport system substrate-binding protein (K02012), *afuB* – iron(III) transport system permease protein (K02011), *afuC* – iron(III) transport system ATP-binding protein (K02010), *HeR* – Heliorhodopsin (PF18761), an 11-subunit respiratory complex I operon (*nuoA, B, C, D, H, I, J, K, L, M, N*), *NDUFAF7* – NADH dehydrogenase [ubiquinone] 1 alpha subcomplex assembly factor 7 (K18164/PF02636), *htpX* – heat shock protein (K03799), *pspE* – phage shock protein E (K03972) containing a *Rhodanese* (PF00581) domain and *adenosine_kinase* (cd01168) (Note: full-length annotated contigs deposited in Figshare). In summary, HeRs from C11 are always preceded by genes encoding a complete ABC-type ferric iron uptake system and followed by an operon encoding a 11-subunit, “ancestral”-type (Moparthi and Hägerhäll 2011) respiratory complex I and by accessory components required for correct assembly and function of this complex (*NDUFAF7, htpX, pspE*) (Zurita Rendón et al. 2014; Pagani and Galante 1983; Alexander and Volini 1987; Sakoh, Ito, and Akiyama 2005). The last conserved gene encodes a pfkB family adenosine kinase (cd01168), a key purine salvage enzyme that phosphorylates adenosine to generate adenosine monophosphate (AMP) (Long, Escuyer, and Parker 2003).

The presence of HeRs within the same transcriptional unit as above mentioned energy metabolism components could indicate them as modulators or even light-induced sensory “switches” of such processes, a mechanism perhaps similar to cryptochrome-driven metabolic synchronization with substrate availability described in other Actinobacteria (Maresca et al. 2019). Notably, beside the 11-subunit complex I, that lacks the NADH dehydrogenase module (subunits nuoE, nuoF, nuoG) (Moparthi and Hägerhäll 2011), *Ca. Actinomarina* (TMED189) genomes also encode the full-sized 14-subunit variant of respiratory complex I in close proximity to the first (for example in GCA-902516125.1). Curiously, the association of HeRs with complex I genes is reminiscent of that between the transmembrane, sensory, EAL-domain containing protein seen in *Bacillus cereus* located upstream of a similar 11-subunit complex I operon (Moparthi and Hägerhäll 2011).

The last cluster featuring a significant number of Actinobacteria HeRs (n = 12) is C17. In this CPR-dominated cluster, Actinobacteria HeRs form a well-defined group sharing a common ancestor with a small CPR sub-cluster. While gene context appears conserved within these Actinobacteria, this does not apply to the putative “sister” CPR sub-cluster.

Clusters C8-C10 share a common ancestor with C11 and include HeRs encoded largely in strict or facultative anaerobic prokaryotes recovered from sediments (including activated sludge). Notably, the phylogenetic tree (iTOL link above) shows two Asgardarchaeota (class Heimdallarchaeia) HeRs branching with very high support (SH-test/UFBoot = 97.2/98) as a sister clade to all C8-C10 members, after the split with the common ancestor shared with C11. Despite the high support for the Asgardarchaeota HeR split, defining a credible cluster will require including additional sequences once more genomes become available. The basal, C10 cluster, is comprised of Archaea-derived HeRs from phyla Crenarchaeota (class Bathyarchaeia) and Euryarchaeota (class Methanobacteria). Clusters C8 and mainly C9 include the MORN-HeR encoding Firmicutes as well as a few Chloroflexota (class Anaerolineae) HeRs.

C12-C16 form a separate super-cluster showing moderate-to-low support for internal branching patterns and include mostly, although not exclusively, anaerobic Archaea (C12 and C14, with the notable exception of aerobic Halobacterota within C12) and anaerobic Chloroflexota (classes Anaerolineae – C13, C15 and Dehalococcoidia – C16). Notably, C16 includes one Thermoplasmatota HeR (encoded in contig SRR5506739-C331) with a zinc-finger extension (Znf-HeR) at the N-terminus. C18 is the most basal cluster with confidently assigned taxonomy. It includes exclusively aerobic Chloroflexota members of the Ellin6529 lineage.

Although no consensus taxonomy could be determined for members of C19 – the first branching group after the split with proteorhodopsins, the abundance of “eukaryotic signature proteins” (ESP) (e.g. Arf, Roc, Rab, etc.) points towards either a eukaryotic origin or unidentified, ESP-rich archaea (Hartman and Fedorov 2002; Dong, Wen, and Tian 2007).

An extended phylogenetic tree of HeRs, including additional dereplicated sequences identified and retrieved from UniProtKB and GTDB, is available in FigShare (see Supp. Methods – Phylogenetic tree of HeRs).

## Supplementary Methods

### Re-assembling of HeR-encoding Spirochaeta

The unexpected detection of HeR in a previously published *Spirochaeta* (diderm organism) genome prompted further investigation. The original Illumina short-read dataset SRX2623364 was downloaded from NCBI SRA (Sequence Read Archive) and preprocessed by using a combination of tools provided by the BBMap project (https://sourceforge.net/projects/bbmap/). This involved removing poor-quality reads with bbduk.sh (qtrim = rl, trimq = 1 8), identifying phiX and p-Fosil2 control reads (k = 21) and removing Illumina sequencing adapters (k = 21). Further, *de novo* assembly of preprocessed paired-end reads was done by Megahit v1.2.9 (D. Li et al. 2016) with k-mer list: 29, 39, 49, 59, 69, 79, 89, 99, 109, 119, 127, and with default parameters. A total of 886 contigs with an average length of 4.98 kbp were produced.

Protein-coding genes were predicted by Prodigal (Hyatt et al. 2010) and taxonomically classified by scanning with MMSeqs2 against the GTDB database. A *Spirochaeta* contig (length = 304,032 bp) was identified and scanned for the presence of HeR against the PFAM database. Both taxonomy (*Spirochaeta*) and HeR presence were consistent with previously published results.

### Phylogenetic tree of HeRs

An extensive collection of predicted HeR amino acid sequences (n = 4,108) was generated from: 1) all HeR sequences available in UniProtKB (n = 502), 2) HeR identified within dereplicated contigs (using cd-hit-est -c 0.95 -aS 0.95) assembled from publicly available metagenomes and metatranscriptomes (n = 3,145), 3) HeR identified within dereplicated high-quality MAGs included in the GTDB database (n = 455) and 4) HeR assembled from the strand-specific metatranscriptomic dataset generated in this study (n = 6). This collection was simplified by keeping only representative sequences (n = 1,669) chosen following clustering with MMSeqs2 (Steinegger and Söding 2017) at 90% sequence identity and 90% coverage (mode: easy-cluster; -c 0.90; --min-seq-id 0.90). Representative HeR sequences were aligned together with 30 selected proteorhodopsins serving as outgroup by using PASTA (Mirarab et al. 2015) (resulting alignment with 1,699 sequences, 2,486 columns, 2,264 distinct patterns, 1,091 parsimony-informative sites, 698 singleton sites, 697 constant sites). A Maximum Likelihood (ML) phylogenetic tree was constructed with IQ-TREE2 (Minh et al. 2020) (1,000 iterations for ultrafast bootstrapping (Hoang et al. 2018) and SH testing, respectively; best model chosen by ModelFinder (Kalyaanamoorthy et al. 2017): LG+G4; additional parameters recommended for short sequence alignments -nstop 500 -pers 0.2). The generated tree was annotated to include labels containing: HeR-encoding contig name, habitat of origin and consensus GTDB taxonomic classification (if available). Data including alignment and the annotated phylogenetic tree are deposited in FigShare (https://figshare.com/s/7bb42426f2ad5e891fec).

### Phylogenetic tree of HeRs with gene context

A simplified depiction of HeR genomic context across representative taxonomic groups harbouring such genes was constructed by merging HeR phylogenetic information with available HeR gene neighbourhood data (Supplementary Figure 4). For this purpose, we established a collection of representative, dereplicated HeR-encoding contigs (n = 872) of at least 5 kb and with clear consensus taxonomy from two sources: 1) HeR-encoding contigs assembled from publicly available metagenomes and metatranscriptomes (n = 3,145) and 2) HeR contigs assembled from the strand-specific metatranscriptomic dataset generated in this study (n = 6). Dereplication of contigs was previously achieved using cd-hit-est (W. Li and Godzik 2006) with identity cutoffs of 95% and coverage of 95% (-c 0.95 -aS 0.95).

### Phylogenetic tree building

Curated HeR sequences recovered from selected contigs were aligned together with 30 proteorhodopsins serving as outgroup by using PASTA (Mirarab et al. 2015) (resulting alignment with 902 sequences with 957 columns, 895 distinct patterns, 550 parsimony-informative sites, 198 singleton sites, 209 constant sites). A Maximum Likelihood (ML) phylogenetic tree was constructed with IQ-TREE2 (Minh et al. 2020) (1,000 iterations for ultrafast bootstrapping (Hoang et al. 2018) and SH testing, respectively; best model chosen by ModelFinder (Kalyaanamoorthy et al. 2017): LG + I+G4; additional parameters recommended for short sequence alignments -nstop 500 -pers 0.2).

### Reconstruction of HeR gene neighborhoods

Coding sequences were predicted *de novo* using Prodigal (Hyatt et al. 2010) in metagenomic (-p meta) mode. Protein domains were annotated by scanning predicted coding sequences against the PFAM (Protein Families) database using the publicly available perl script pfam_scan.pl. Predicted protein sequences from all contigs were clustered together using the MMSeqs2 easy-cluster workflow with 50% identity and 80% coverage cutoffs (--min-seq-id 0.5 -c 0.8 -e1 e-3 -- cluster-reassign) and a minimum of 2 sequences per cluster. Clusters were sorted according to their size (i.e. number of sequences) and colour codes were assigned to the top largest 63. All clusters containing HeRs were assigned matching colours. HeR gene neighbourhood data was combined with the reconstructed HeR phylogenetic tree in iTOL (Letunic and Bork 2016) (https://itol.embl.de/). To facilitate visual interpretation, a number of adjustments and rules were applied: 1) all contigs were oriented according to the sense (+) of encoded HeRs, 2) contigs were centered on HeR genes with a maximum of 10 neighbouring genes depicted up- and downstream, 3) information regarding gene lengths was not included, all of them being shown as equally sized rectangles, 4) homologous genes (i.e. members of the same MMSeqs2 defined cluster) share matching colours within each phylogenetically defined cluster, 5) all HeRs are coloured the same across all phylogenetic clusters, 6) grey rectangles indicate genes with few homologues and/or singletons, 7) taxonomy, habitat information and phylogenetic clusters are colour coded on independent strips.

### MORN-HeR phylogenomic tree

A maximum-likelihood (ML) phylogenomic tree was constructed for Firmicutes MAGs encoding MORN-HeR protein domain fusions along with other representatives of this phylum that assumably lack such genes. The established collection of (n = 68) MAGs and reference genomes (Supplementary Table S15) was scanned by hmmsearch against a previously published list of (n = 120) conserved protein marker HMMs (Parks et al. 2018). Four divergent markers (TIGR00442, TIGR00539, TIGR00643, TIGR00717) were identified by scanning with CD-search (e-value < 1e-2) and removed. Curated amino acid sequences for the selected 116 phylogenetic markers were aligned with PRANK (Löytynoja 2014) and resulting alignments were trimmed by BMGE (parameters: -t AA -g 0.5 -b 3 -m BLOSUM30) (Criscuolo and Gribaldo 2010). Individually trimmed alignments were concatenated resulting in a block of 68 sequences with 40,714 columns, 36,088 distinct patterns, 28,978 parsimony informative sites, 3,366 singleton sites and 8,370 constant sites. The concatenated alignment was used with IQ-TREE (v.1.6.12) to construct the ML phylogenomic tree (parameters: 1,000 iterations of ultrafast bootstrapping (Hoang et al. 2018) and SH testing (Minh et al. 2020), respectively; best model (LG + F+R5) chosen by ModelFinder (Kalyaanamoorthy et al. 2017).

### Multiple sequence alignment (MSA) of MORN-HeR proteins

Predicted amino acid sequences containing full-length MORN-MORN-MORN-HeR protein domain fusions (n = 36) were retrieved from Firmicutes MAGs (n = 35) previously used to construct the phylogenomic tree presented in Supplementary Figure 1. All sequences were aligned together using the PSI-Coffee alignment method (Chang et al. 2012) provided by the T-Coffee online server (http://tcoffee.crg.cat) with default parameters. The resulting MSA is presented with annotations in Supplementary Figure 2 while the original alignment file generated by PSI-Coffee is available in FigShare (https://figshare.com/s/7bb42426f2ad5e891fec).

---

**Supplementary Figure 1.**
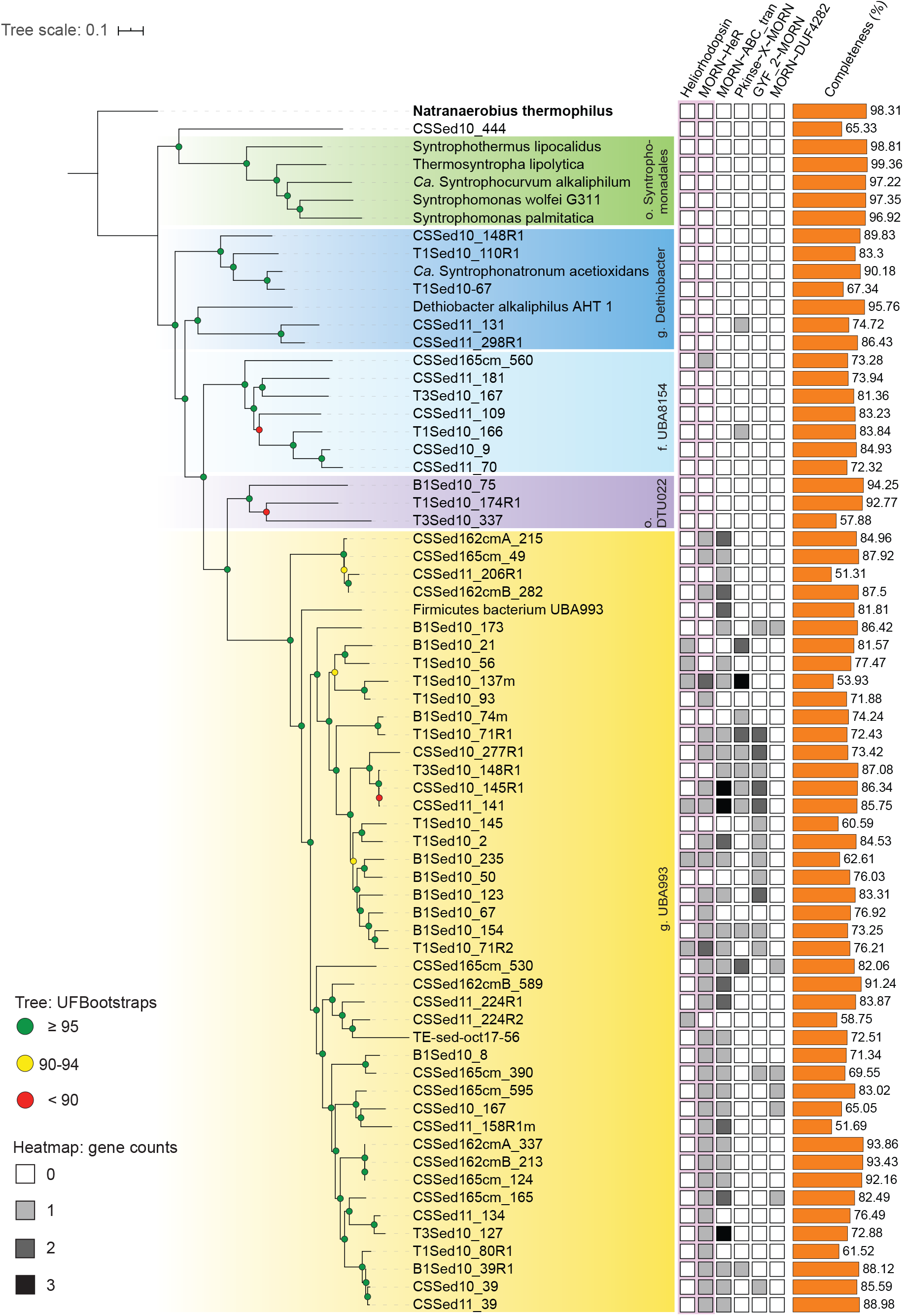
Phylogenomic tree of MORN-Heliorhodopsin (MORN-HeR) encoding MAGs. Green circles indicate high confidence UFBootstrap values (≥ 95). Occurences of genes encoding heliorhodopsins, MORN-heliorhodopsin as well as other MORN-domain fusions are depicted in the adjacent matrix. Genome completeness values are depicted as a histogram (estimated by CheckM). All genomes are members of the *Firmicutes* phylum, with different taxonomic subdivisions highlighted on the tree (taxonomy by GTDBtk). *Natranaerobius thermophilus* was used to root the tree. The majority of genomes included here have been previously used for phylogenomic analyses by Timmers et. al, 2018. Among reference genomes, *Firmicutes bacterium UBA993* and *Ca. Syntrophocurvum alkaliphilum* are new additions.

**Supplementary Figure 2.**
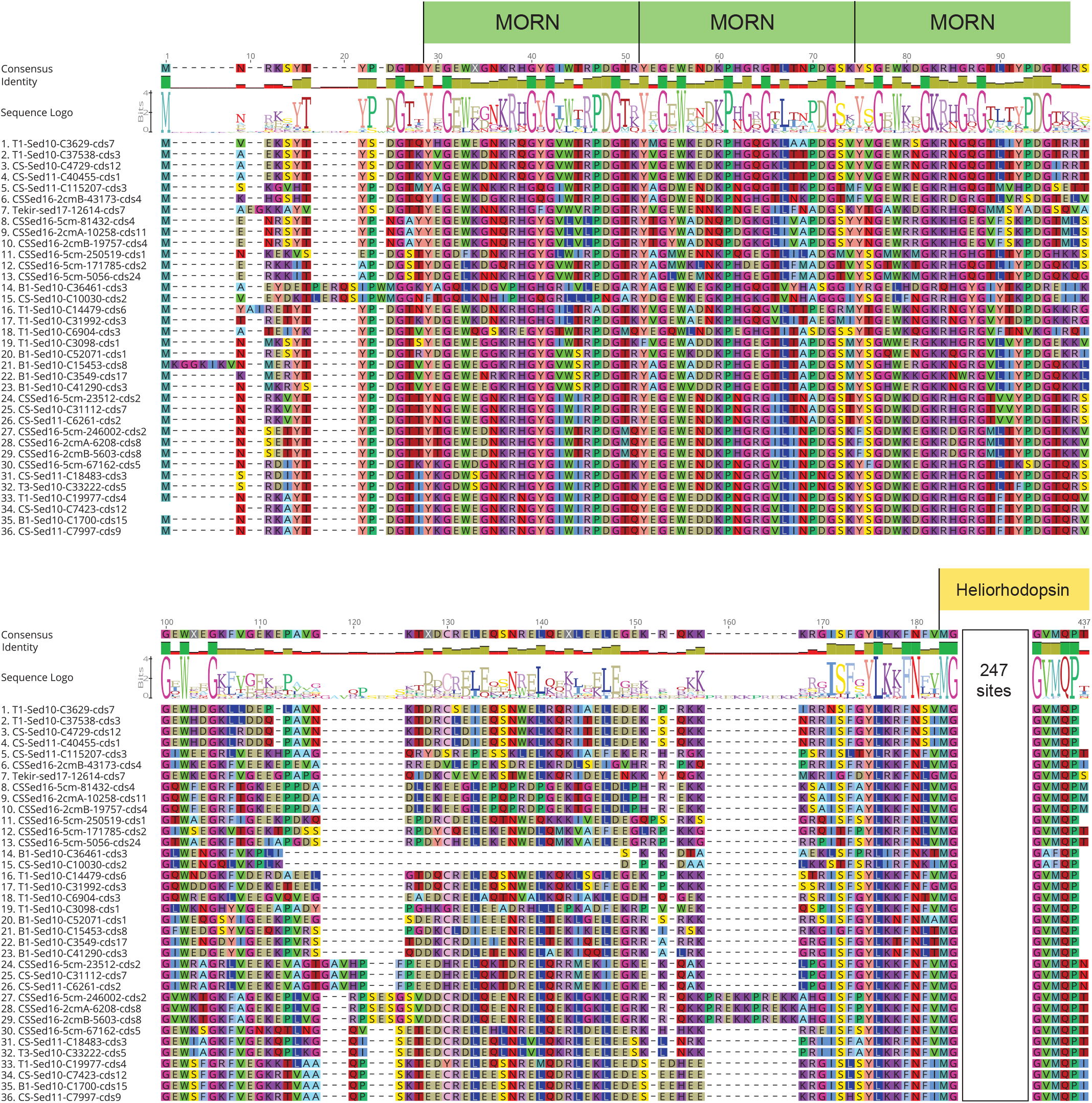
Multiple sequence alignment of MORN-HeR protein domain fusions predicted in Firmicutes MAGs. Each sequence shows 3 consecutive MORN domains (indicated by green rectangles). Aligned HeR domains display a high level of conservation and are truncated for illustration purposes (indicated by yellow rectangle). The original full-length alignment was deposited in FigShare.

**Supplementary Figure 3.**
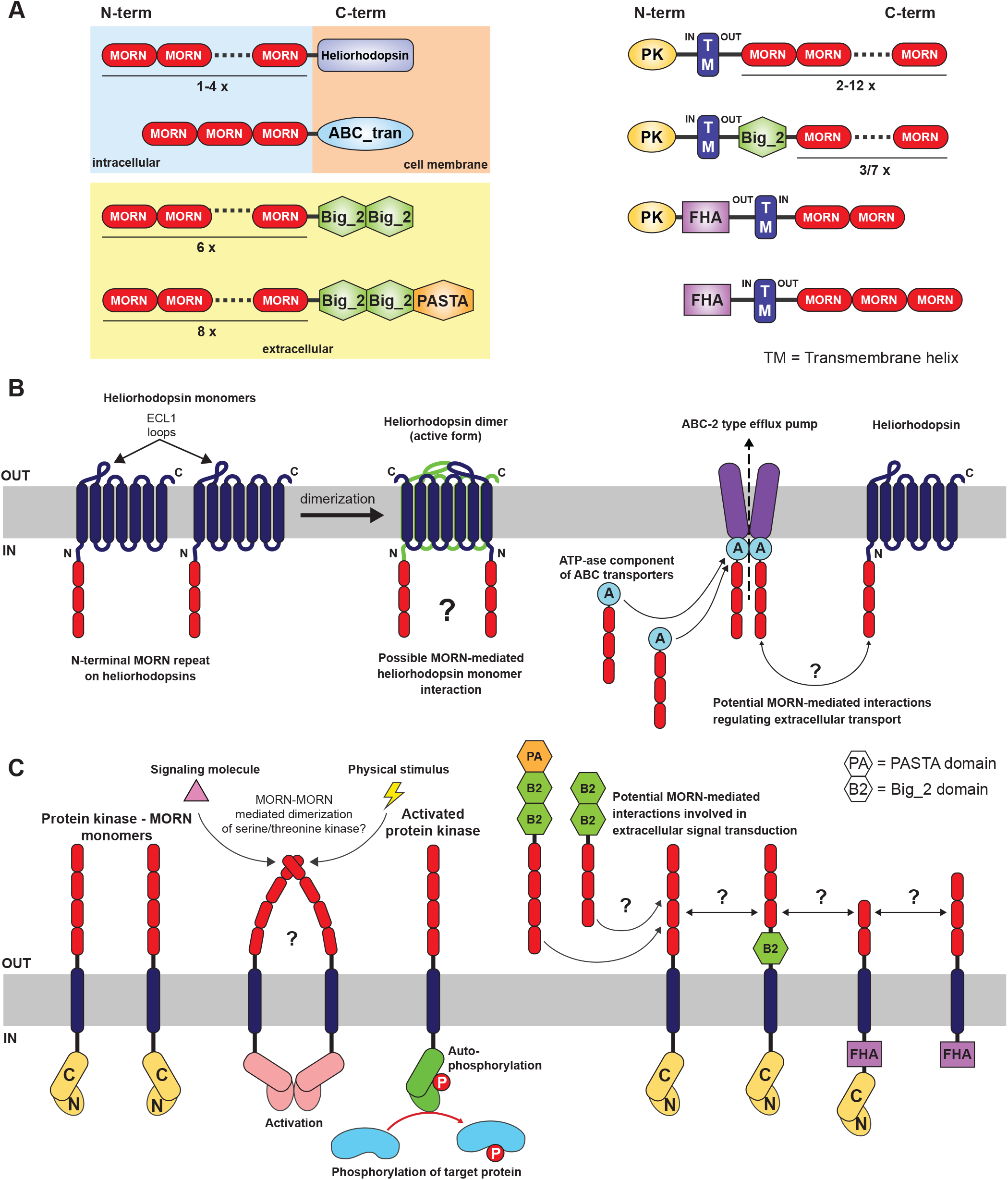
Summary of MORN-repeat proteins predicted from metagenome-assembled genomes (MAGs) of anaerobic Firmicutes. **A**). Schematics of frequently co-occuring MORN-repeat containing proteins in Firmicutes MAGs including MORN-HeR, MORN-ABC transporters – likely involved in drug resistance or Cu^2+^/Na^2+^ ion efflux, periplasmic proteins containing bacterial immunoglobulin-like folds (Big_2, PASTA), MORN-protein kinase fusions where MORN repeats and p-kinase domains are commonly separated by transmembrane α-helices on opposite sides of the cellular membrane and MORN-forkhead associated (FHA) domains. **B**) Potential interactions between MORN-HeR monomers, MORN-ATPase components of ABC transporters and MORN-HeR and ABC transporters mediated or stabilized by the presence of MORN-repeat fusions. **C**) Potential interactions of MORN-Protein-kinases associated with functions such as extracellular signal transduction.

**Supplementary Figure 4.**
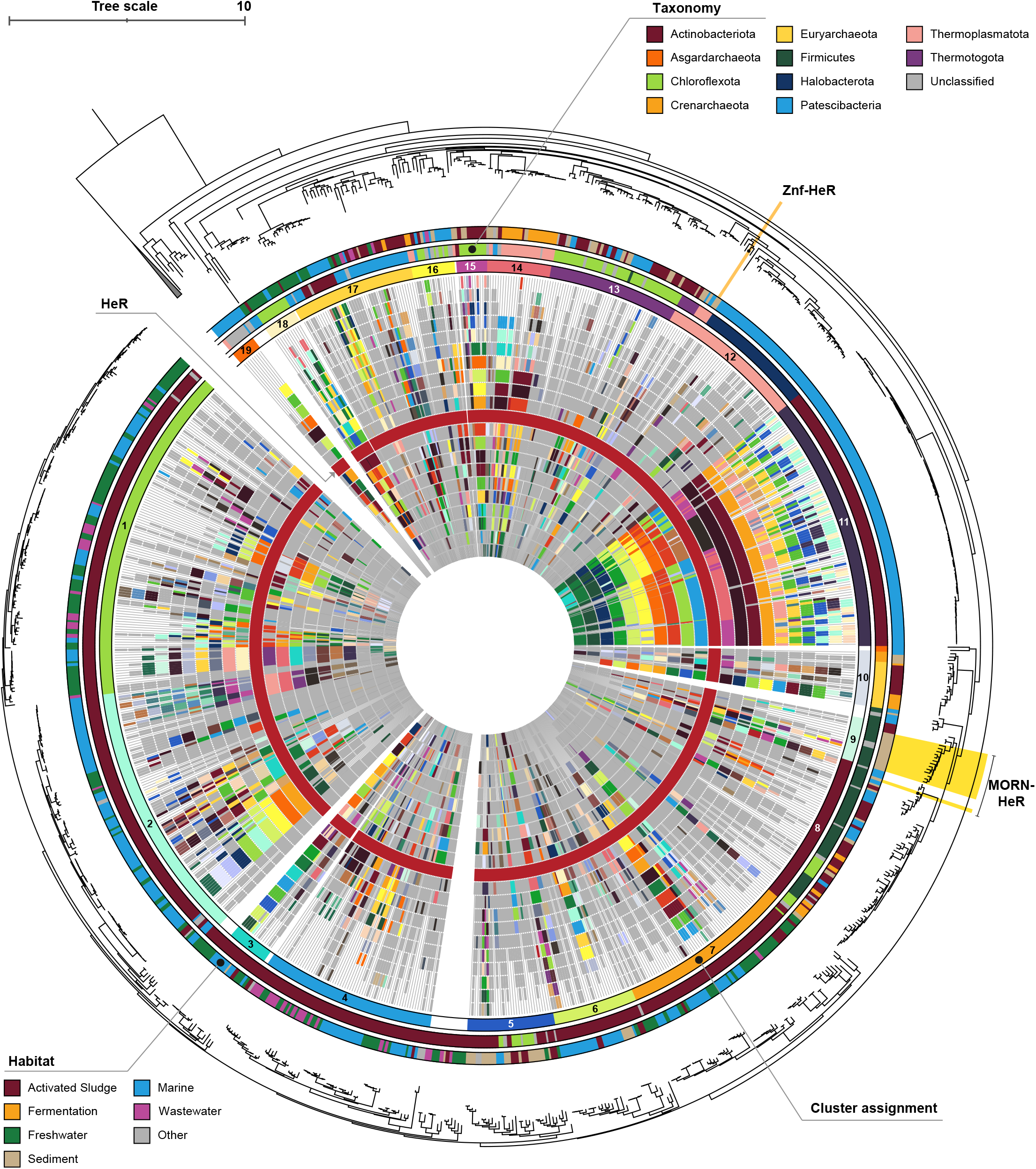
Genomic context of heliorhodopsin (HeR) genes across representative taxonomic groups. The phylogenetic tree was constructed using 872 HeR amino acid sequences and 30 proteorhodopsins used as an outgroup for rooting. Gene neighbourhoods (10 genes up- and downstream) for each HeR were depicted schematically. Abundant homologues are coloured within each defined phylogenetic cluster while less abundant genes and/or singletons are depicted in gray. All contigs were centered and oriented according to encoded HeR (dark red circle). Information regarding relative gene lengths was not included. Particular HeR-protein domain fusions are indicated separately: Znf-HeR and MORN-HeR.

**Supplementary Figure 5.**
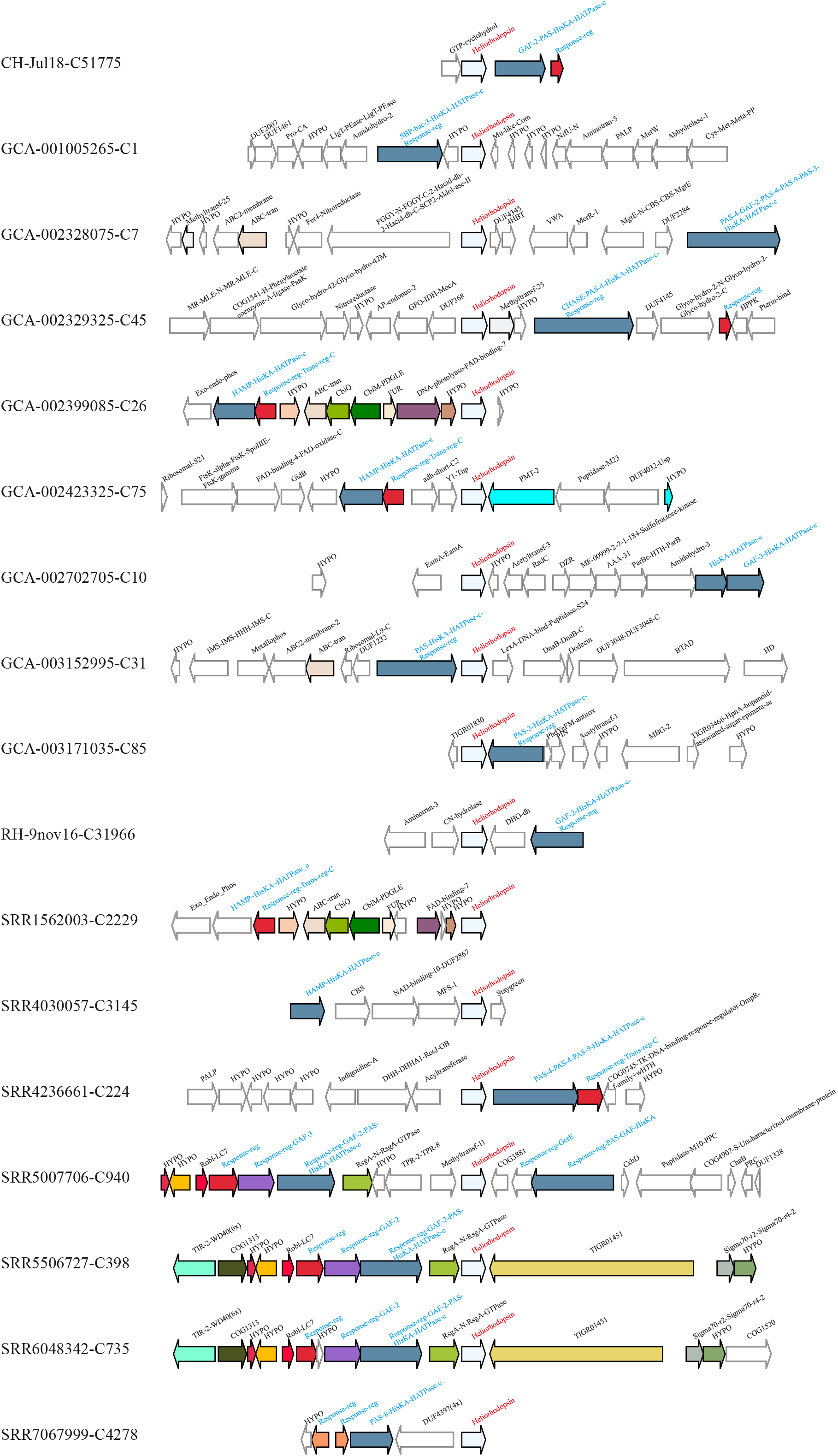
Histidine kinases and related genes (blue labels) in the genomic neighborhood (within 10 kb) of Heliorhodopsins (red labels) originating from the phylum Chloroflexi.

**Supplementary Figure 6.**
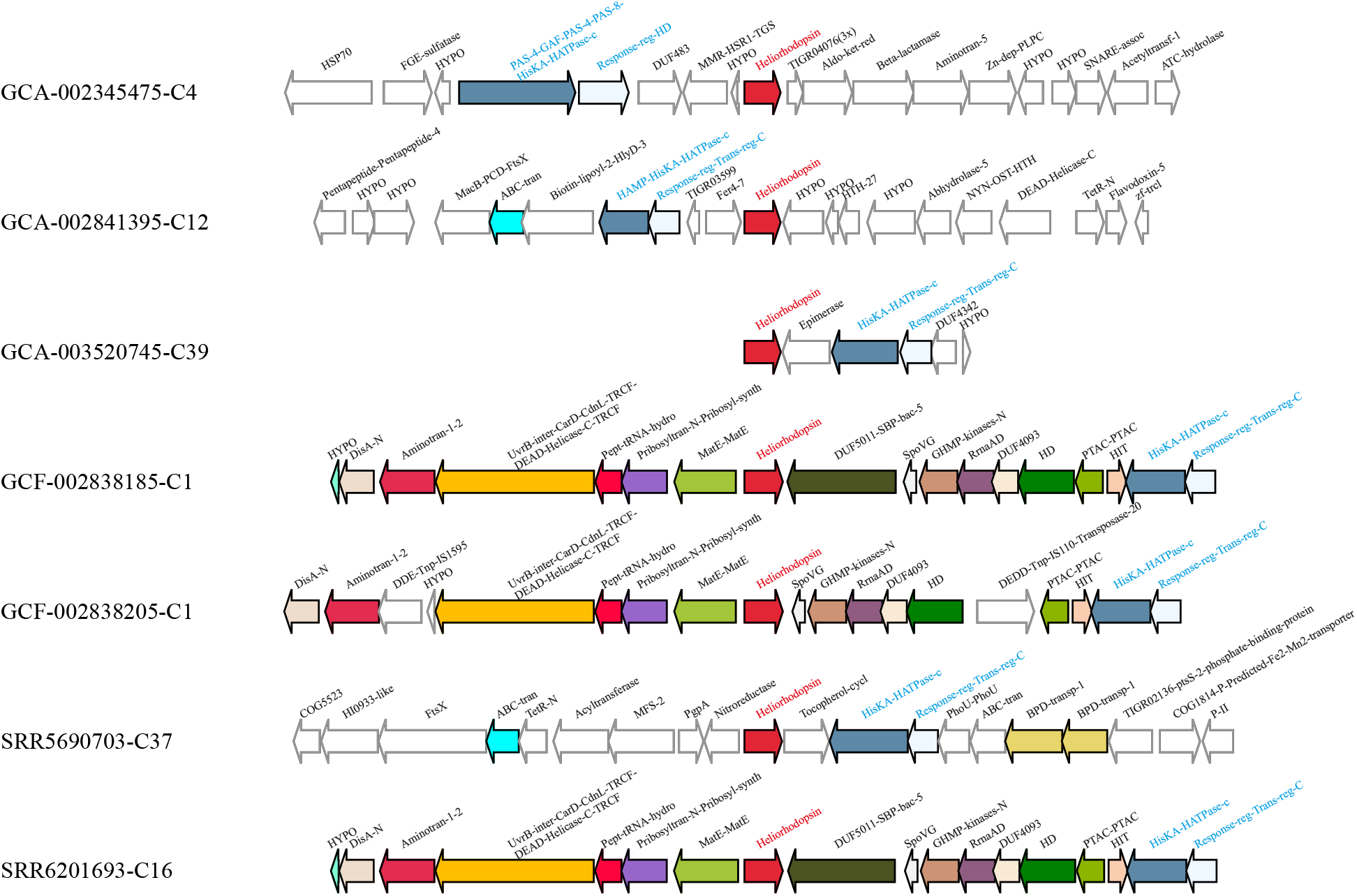
Histidine kinases and related genes (blue labels) in the genomic neighborhood (within 10 kb) of Heliorhodopsins (red labels) originating from the phylum Firmicutes.

**Supplementary Figure 7.**
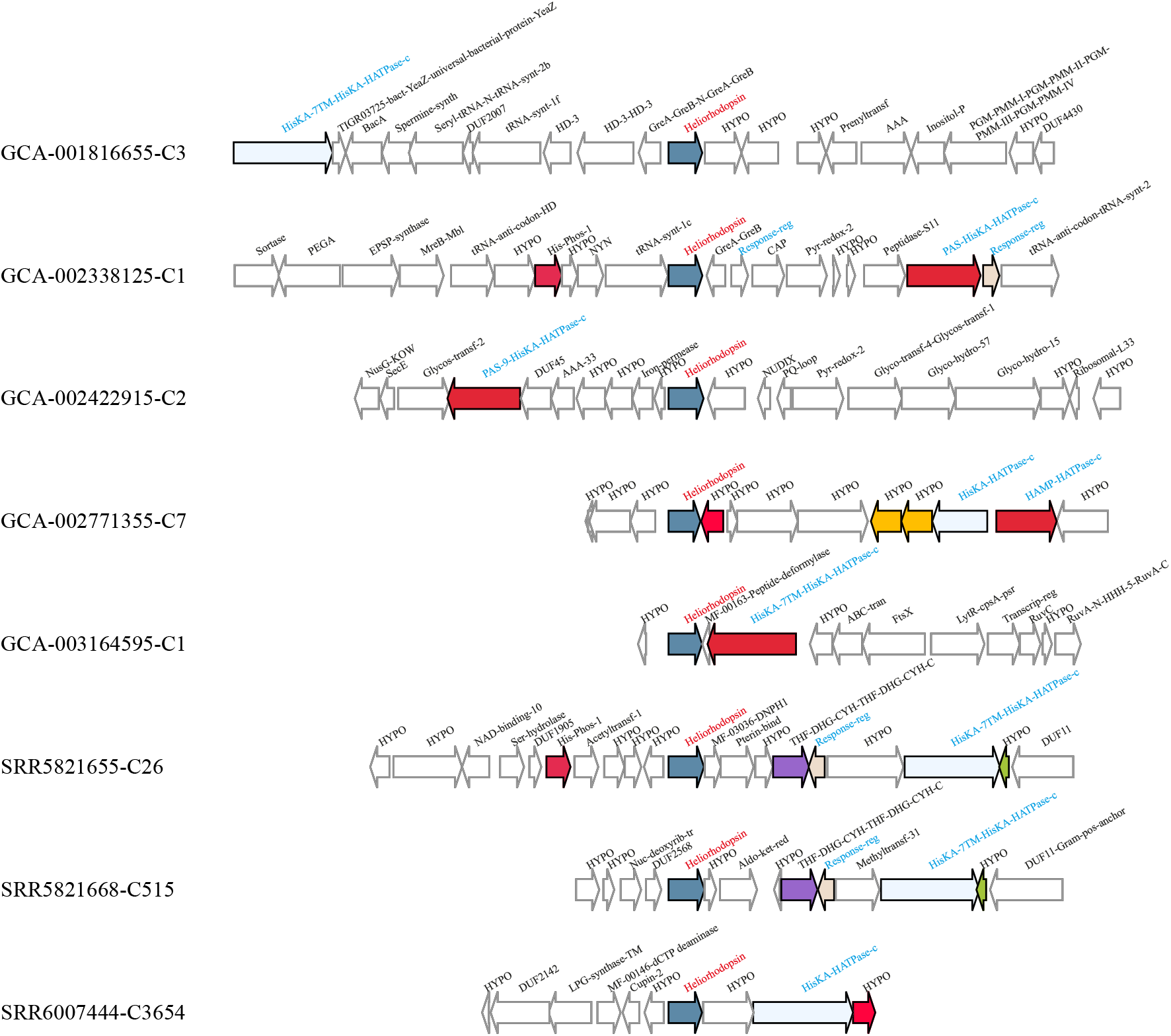
Histidine kinases and related genes (blue labels) in the genomic neighborhood (within 10 kb) of Heliorhodopsins (red labels) originating from the phylum Patescibacteria.

**Supplementary Figure 8.**
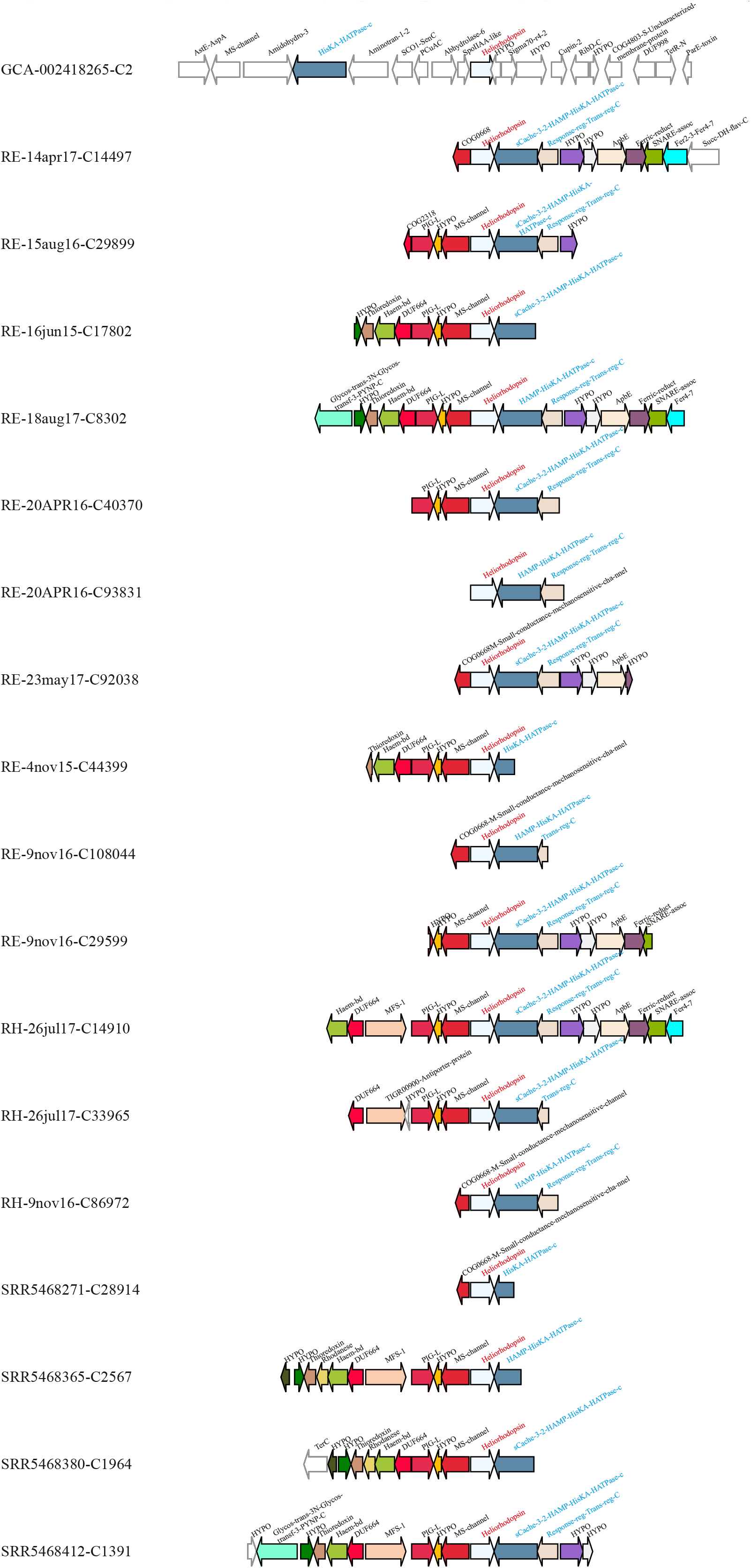
Histidine kinases and related genes (blue labels) in the genomic neighborhood (within 10 kb) of Heliorhodopsins (red labels) originating from the phylum Actinobacteriota, class Acidimicrobiia.

**Supplementary Figure 9.**
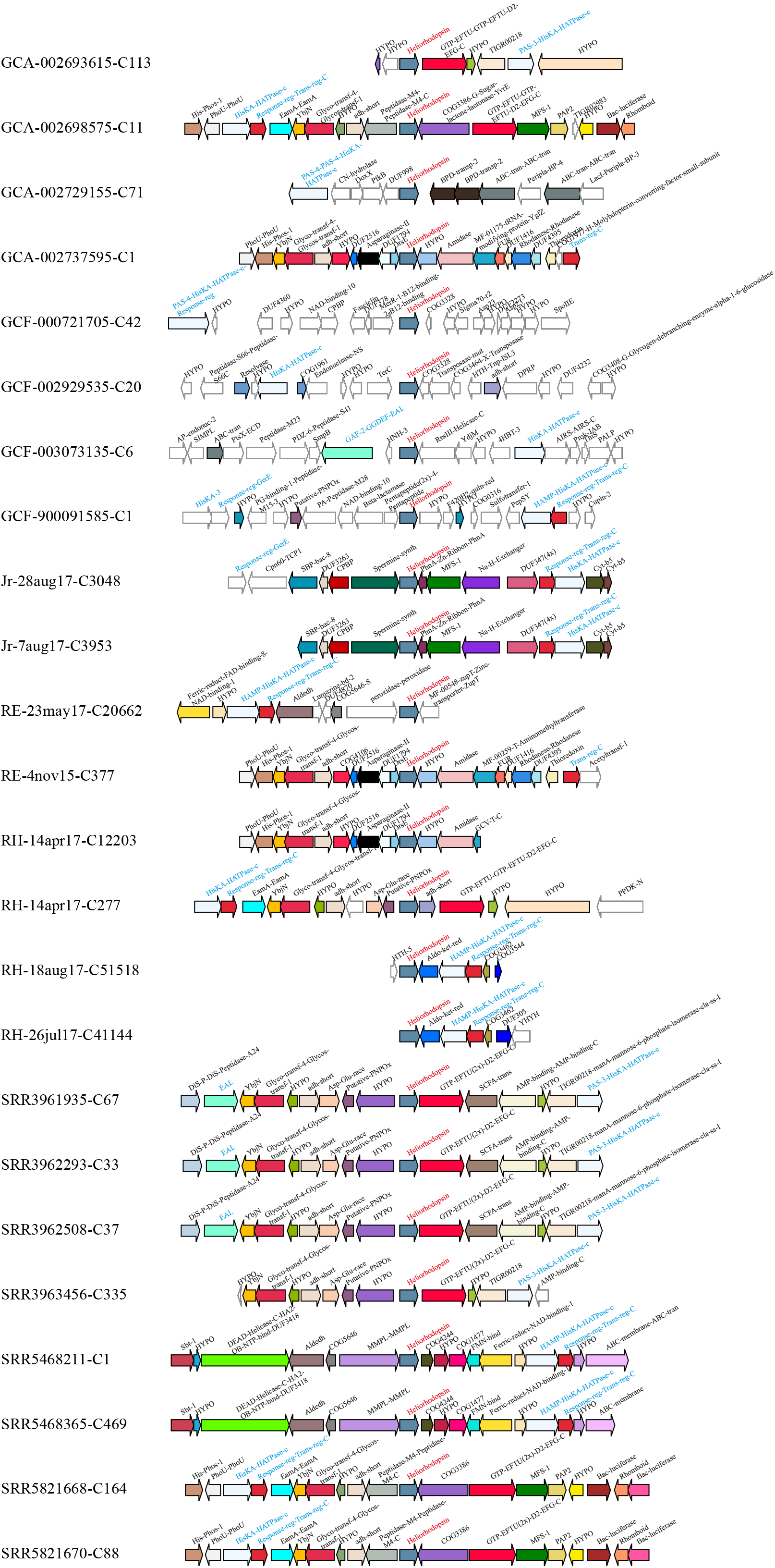
Histidine kinases and related genes (blue labels) in the genomic neighborhood (within 10 kb) of Heliorhodopsins (red labels) originating from the phylum Actinobacteriota, class Actinobacteria.

**Supplementary Figure 10.**
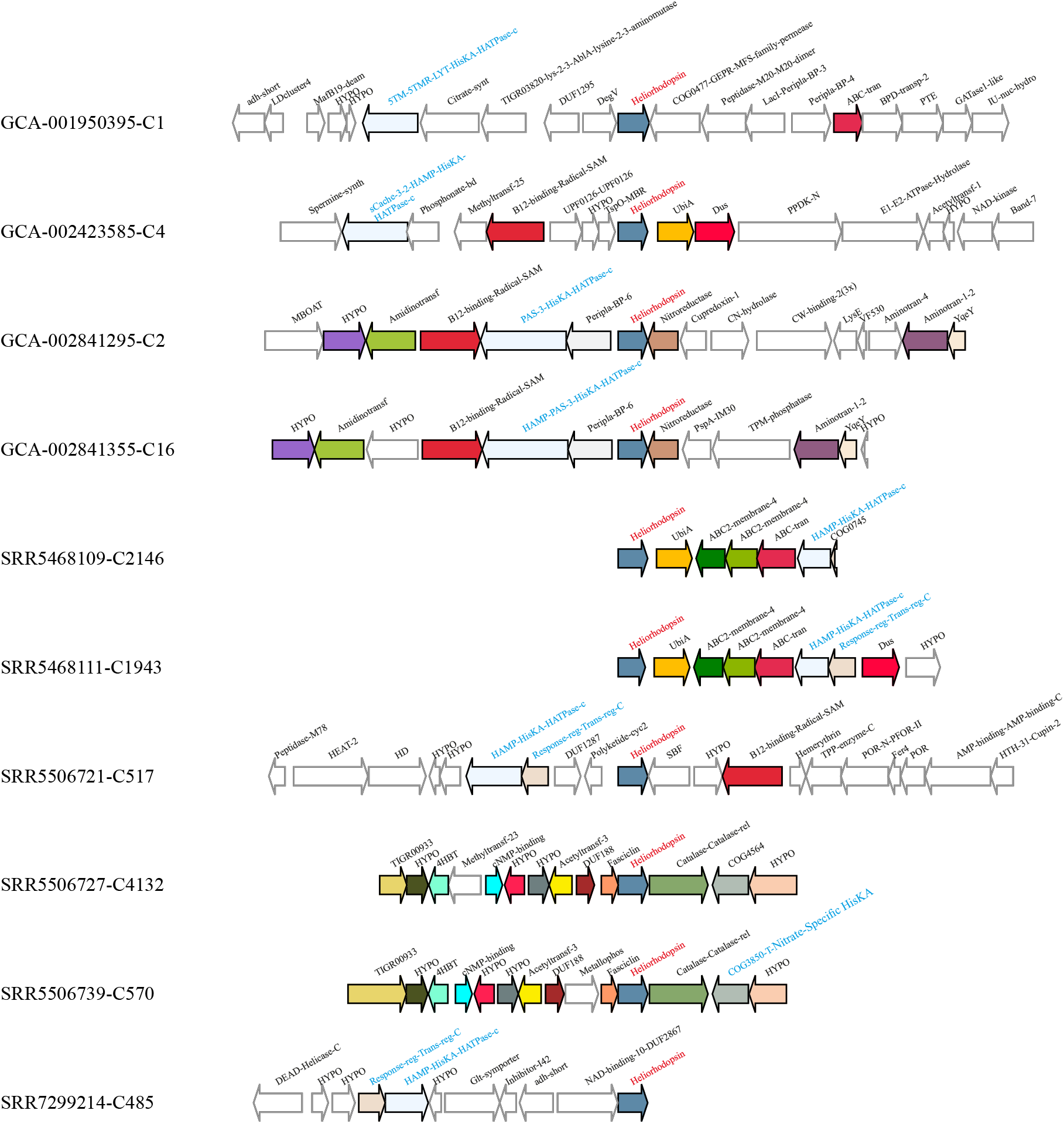
Histidine kinases and related genes (blue labels) in the genomic neighborhood (within 10 kb) of Heliorhodopsins (red labels) originating from the phylum Actinobacteriota, class Coriobac-teriia.

**Supplementary Figure 11.**
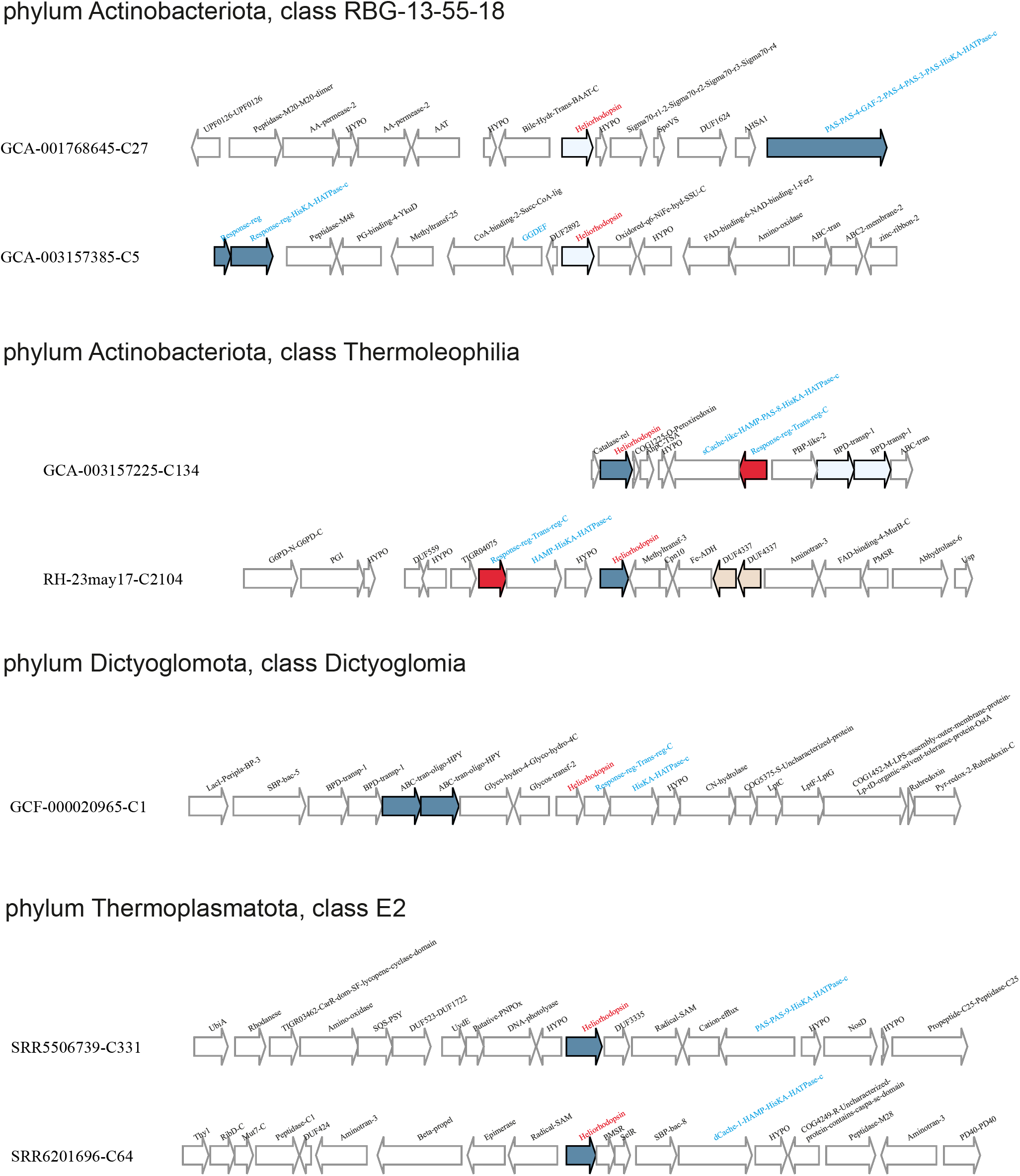
Histidine kinases and related genes (blue labels) in the genomic neighborhood (within 10 kb) of Heliorhodopsins (red labels) originating from the phylum Actinobacteriota (classes RBG-13-55-18 and Thermoleophilia), phylum Dictyoglomota (class Dictyoglomia) and phylum Thermoplas-matota (class E2).

**Supplementary Figure 12.**
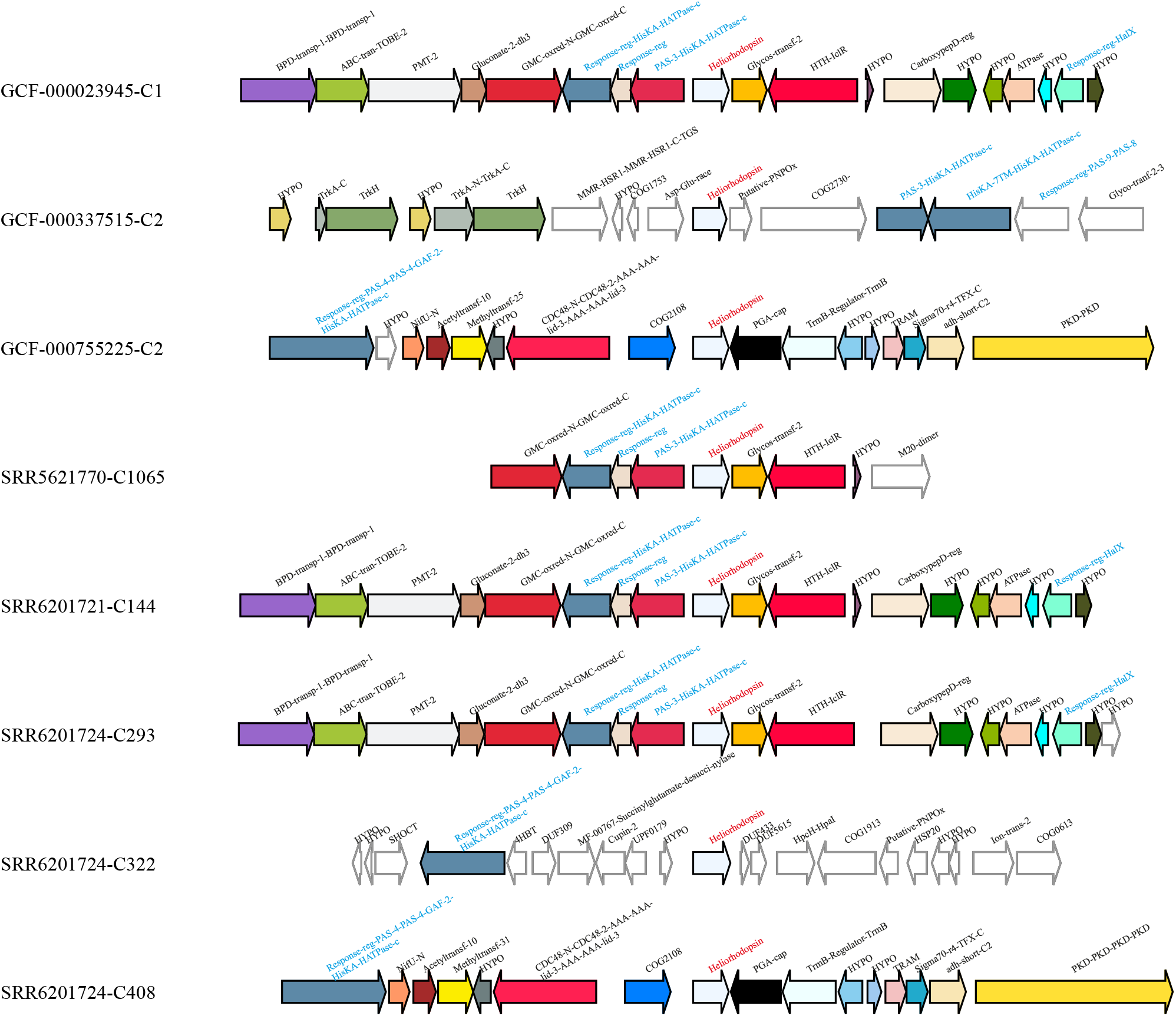
Histidine kinases and related genes (blue labels) in the genomic neighborhood (within 10 kb) of Heliorhodopsins (red labels) originating from the phylum Halobacterota, class Halobacteria.

**Supplementary Figure 13.**
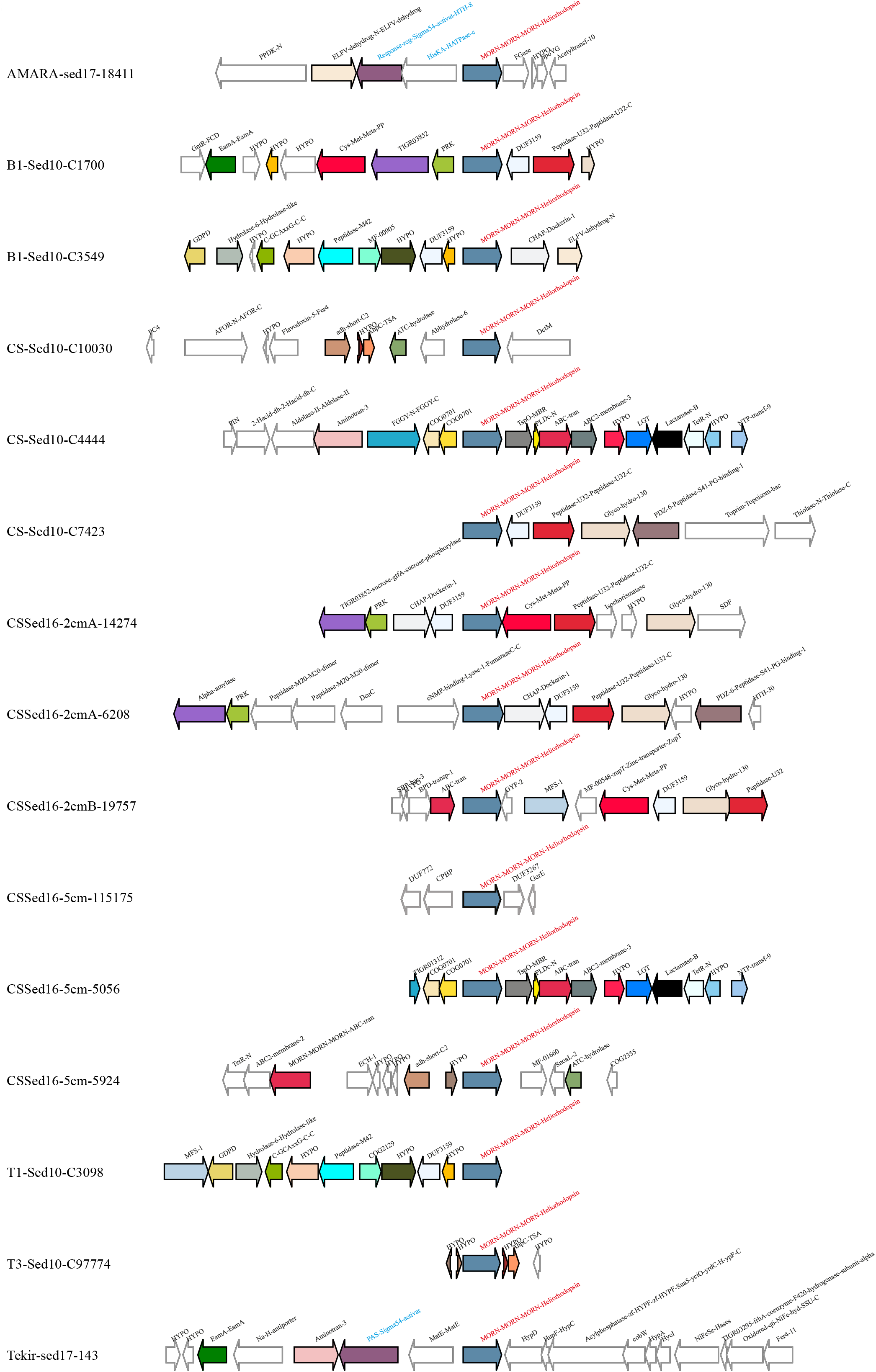
Histidine kinases and related genes (blue labels) in the genomic neighborhood (within 10 kb) of MORN-Heliorhodopsins (red labels) originating from Firmicutes (phylum) MAGs (sediment and brine metagenomes).

**Supplementary Figure 14.**
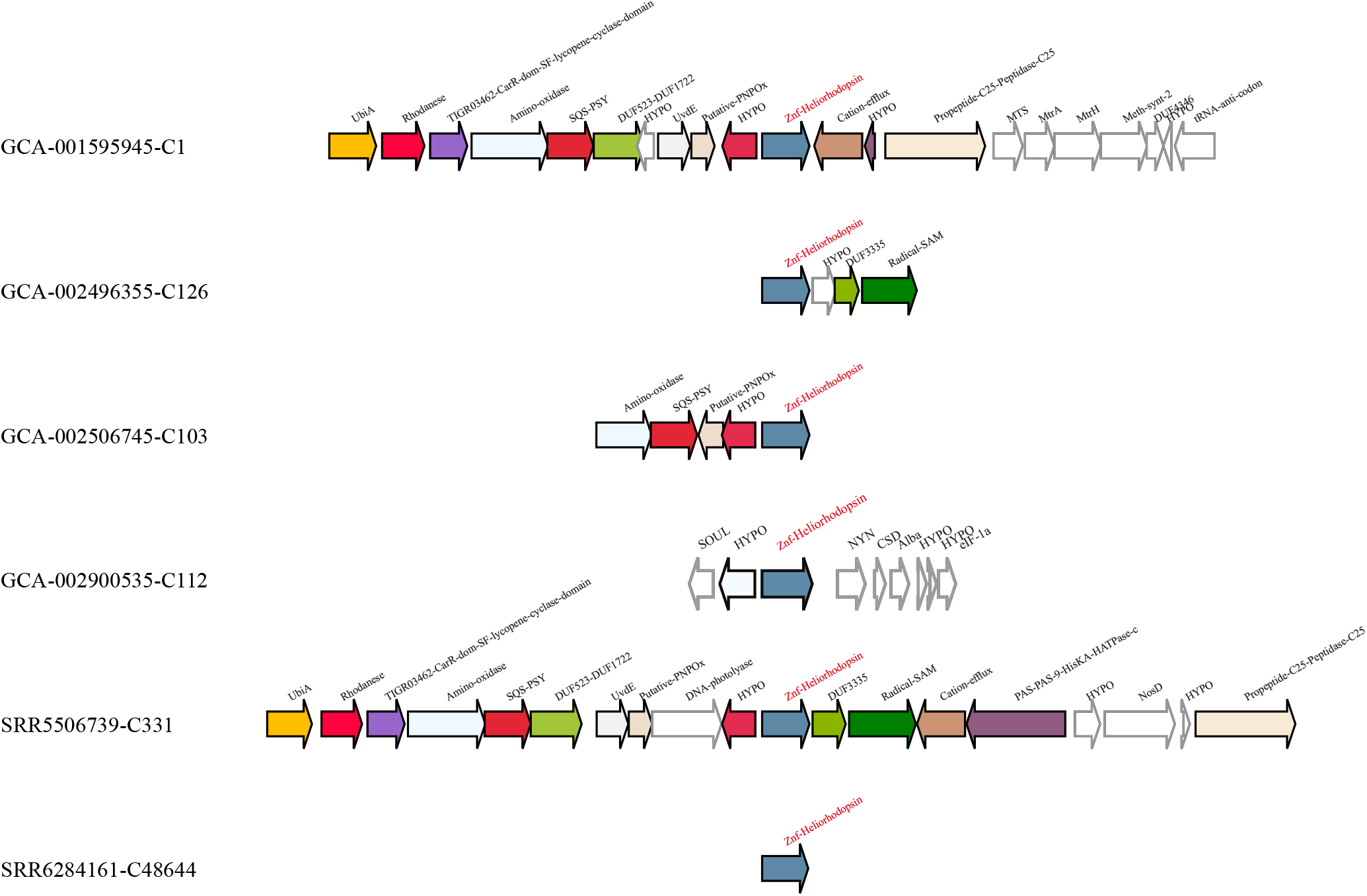
Genomic neighborhood (within 10 kb) of Znf-Heliorhodopsins (red labels) originating from archaeal contigs from the phylum Thermoplasmatales, class E2.

**Supplementary Figure 15.**
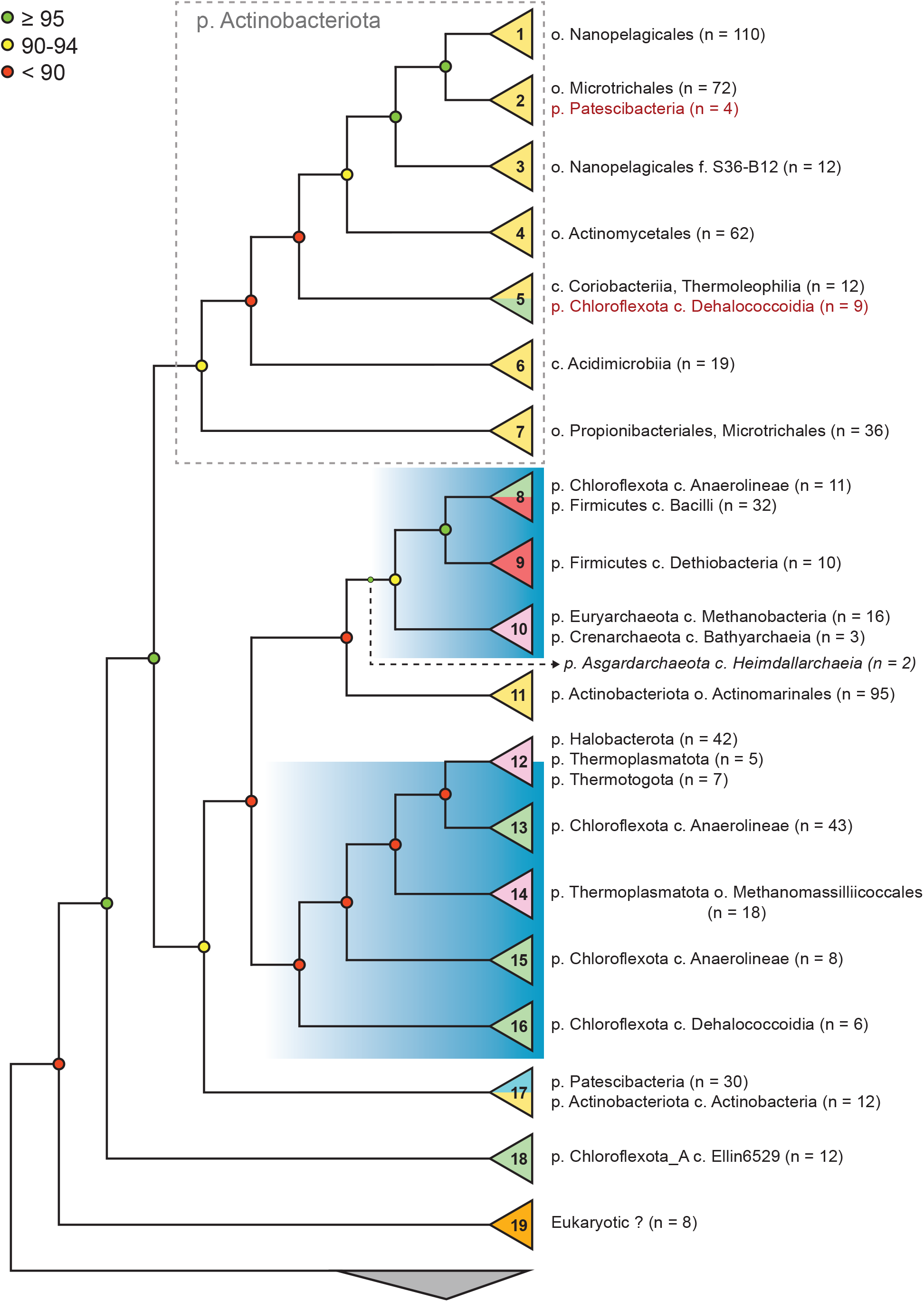
Simplified representation (cladogram) of the HeR phylogenetic tree used for gene context analysis. Cluster numbers (defined in Supp. Fig. 4) are indicated on triangles at the tip of each branch. Actinobacteriota clusters are coloured yellow, Archaea - purple, Chloroflexota - green, Eukaryota - orange, Firmicutes - red, Patescibacteria - blue. Taxonomy and sequence counts are shown-only for representatives of each cluster. The blue rectangles highlight clusters where all or most members are anaerobic organisms. The outgroup (proteorhodopsins; n = 30) is depicted as a gray triangle at the bottom.

